# ZNF143 is a transcriptional regulator of nuclear-encoded mitochondrial genes that acts independently of looping and CTCF

**DOI:** 10.1101/2024.03.08.583864

**Authors:** Mikhail D. Magnitov, Michela Maresca, Noemí Alonso Saiz, Hans Teunissen, Luca Braccioli, Elzo de Wit

## Abstract

Gene expression is orchestrated by transcription factors, which function within the context of a three-dimensional genome. Zinc finger protein 143 (ZNF143/ZFP143) is a transcription factor that has been implicated in both gene activation and chromatin looping. To study the direct consequences of ZNF143/ZFP143 loss, we generated a ZNF143/ZFP143 degron line. Our results show that ZNF143/ZFP143 depletion has no effect on chromatin looping. Systematic analysis of ZNF143/ZFP143 occupancy data revealed that a commonly used antibody cross-reacts with CTCF, leading to its incorrect association with chromatin loops. Nevertheless, ZNF143/ZFP143 specifically activates nuclear-encoded mitochondrial genes and its loss leads to severe mitochondrial dysfunction. Using an *in vitro* embryo model, we find that ZNF143/ZFP143 is an essential regulator of organismal development. Our results establish ZNF143/ZFP143 as a conserved transcriptional regulator of cell proliferation and differentiation by safeguarding mitochondrial activity.

**Highlights:**

1. Acute degradation of ZFP143 leads to rapid and specific loss of gene transcription.
2. Molecular consequences of ZFP143 loss are inconsistent with a role in chromatin looping.
3. ZFP143 is a conserved transcriptional regulator of nuclear-encoded mitochondrial proteins.
4. ZFP143 regulation of mitochondrial homeostasis is critical for multicellular organismal development.

**Graphical Abstract:** 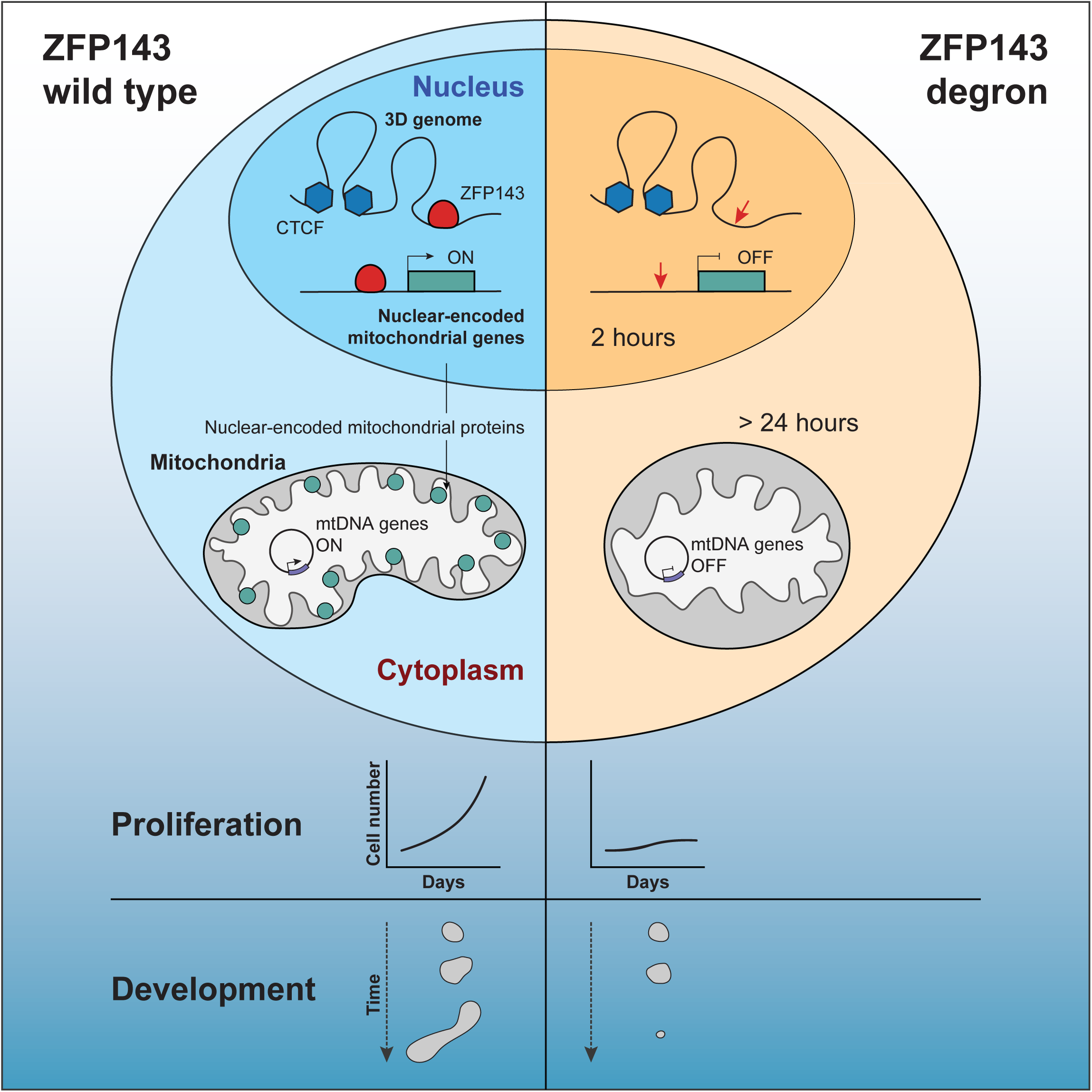

## Introduction

Gene expression is a key biological process that is regulated by transcription factors (TFs), which can recognise and bind specific regulatory DNA sequences^1^. Some TFs can act by binding directly to gene promoters, while others can modulate expression by targeting distal enhancers, which can be located tens or even hundreds of kilobases away from their target promoters^2–4^.

Zinc finger protein 143 (ZNF143/ZFP143, referring to the human/mouse proteins, respectively) is a sequence-specific DNA-binding transcription factor. It is an orthologue of selenocysteine tRNA gene transcription activating factor (Staf) originally identified in *Xenopus laevis*^5–7^. Soon after its discovery, Staf was described as a transcriptional activator of snRNA genes transcribed by RNA polymerases II and III^8^ and was later shown to also be capable of activating transcription from an mRNA promoter^7,9^. Transcriptional activation by Staf is mediated by recognition of its consensus DNA binding motif, the Staf binding site (SBS)^5,8^.

Genome-wide analyses in human and mouse cells revealed that the SBS motifs are present in thousands of promoters and are occupied by ZNF143^10–12^. Subsequent experiments confirmed that ZNF143 binding to promoter regions directly regulates the expression of its target genes^12,13^. ZNF143 is expressed in a wide range of tissues^14^ and was identified as an ‘essential’ gene in a haploid genetic screen^15^ and a ‘common essential’ gene in DepMap^16^. Given its established role as a transcriptional regulator, ZNF143 has been implicated in many biological, pathological and developmental processes, such as proliferation and cell cycle regulation^12,17–20^, cancer cell survival^18,20–22^, embryonic development^23–26^ and many others^27,28^. However, how the regulatory functions of ZNF143 converge to give rise to these diverse cellular functions remains to be elucidated.

Recent studies have suggested that ZNF143 is also involved in spatial genome organisation. Two key players in the organisation of chromosomes are the cohesin complex, which is capable of binding to and extruding DNA to create chromatin loops, and CTCF, which can create barriers to the extrusion process by blocking cohesin^29^. Multiple studies reported that ZNF143 colocalises with CTCF^30–39^ and participates in chromatin looping^33–47^. Deletion or silencing of ZNF143 resulted in a weakening of a subset of chromatin interactions, particularly those centred on promoter and enhancer regions^34,38,39,46,47^, raising the possibility that ZNF143 may function in conjunction with CTCF to establish enhancer-promoter loops. However, these studies were performed in a non-acute depletion setting, making it difficult to disentangle the direct and indirect effects of ZNF143 loss on chromatin structure.

In this work, we aimed to understand the direct role of ZNF143 in chromatin looping and its interplay with transcriptional regulation of target genes. To study this, we used an acute protein degradation system to rapidly deplete ZFP143 from mouse embryonic stem cells (mESCs). By measuring chromatin occupancy and spatial chromatin interactions, we found that ZFP143 does not play a role in chromatin looping. Systematic re-analysis of publicly available ZNF143/ZFP143 binding datasets revealed a strong antibody bias, which we propose led to the detection of co-localisation with CTCF and subsequent association with chromatin looping. Using nascent transcriptome profiling, we find that ZFP143 acts as a strong transcriptional regulator of nuclear-encoded mitochondrial proteins, in particular genes encoding proteins of the mitochondrial ribosome and oxidative phosphorylation complexes. Our analyses establish ZNF143/ZFP143 as a conserved transcriptional regulator of cellular proliferation and differentiation that acts independently of chromatin looping and CTCF.

## Results

### Establishment of ZFP143 degron cell line

To investigate the direct impact of ZFP143 loss on chromatin structure and gene expression, we utilised the degradation tag (dTAG) system to rapidly and acutely deplete ZFP143 from mESCs^48^. Using CRISPR-mediated genome editing, we endogenously tagged the C-terminal domain of ZFP143 with the FKBP12^F36V^ domain, 2xHA tag and eGFP (Figure 1A, Table S1). We characterised the kinetics of ZFP143 loss by treating mESCs with the dTAG-V1 small molecule, which targets the fusion protein to the von Hippel-Lindau (VHL) E3 ubiquitin ligase, ultimately resulting in rapid protein degradation^49^. Immunoblotting showed that ZFP143 could be depleted from the established mESCs within 2 hours of dTAG-V1 treatment (Figure 1B, 1C). To further validate the depletion of ZFP143, we performed ChIP-seq using an antibody against the introduced HA tag. We found that ZFP143 was no longer detectable on chromatin after 2 hours of dTAG-V1 treatment (Figure 1D), suggesting efficient depletion of the target protein.

**Figure 1.**
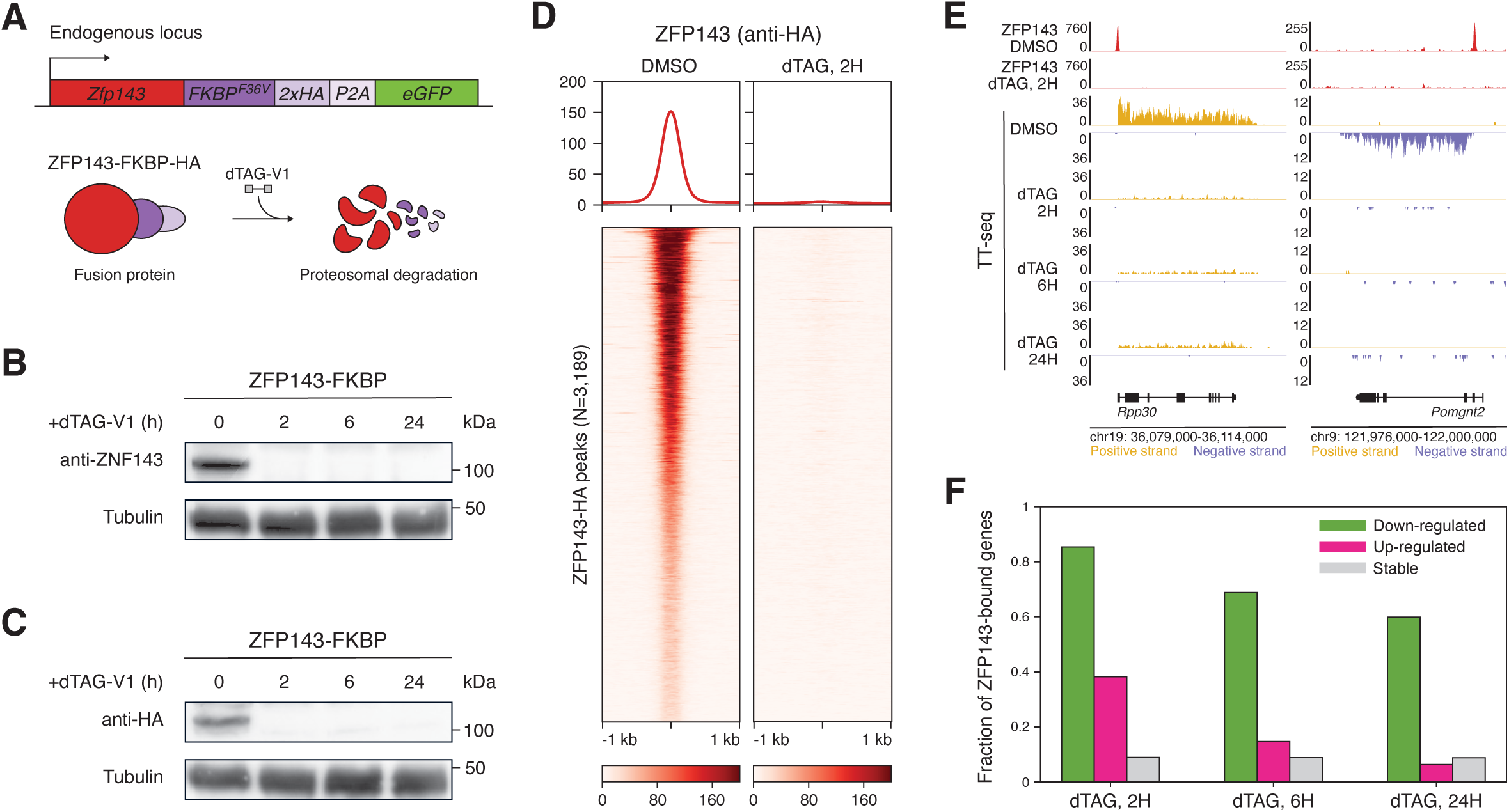
Generation and validation of a system for rapid depletion of ZFP143 in mESCs. **(A)** Schematic representation of the dTAG degron system. The endogenous *Zfp143* locus is modified to create a fusion with the FKBP12F36V-2xHA-P2A-eGFP cassette. The FKBP-tagged ZFP143 protein can be degraded through the VHL protein degradation pathway upon addition of the dTAG-V1 small molecule. **(B)** Western blot of ZFP143-FKBP degron cells with an antibody against the endogenous ZFP143. ZFP143 protein abundance is visualised upon addition of dTAG-V1 for the indicated times. ɑ-Tubulin was used as loading control. **(C)** Western blot of ZFP143-FKBP degron cells with an antibody against the HA tag. ZFP143-HA protein abundance is visualised upon addition of dTAG-V1 for the indicated times. ɑ-Tubulin was used as loading control. **(D)** Tornado plots showing ZFP143-HA ChIP-seq signal centred at ZFP143-HA peaks in DMSO and 2 hours dTAG-V1 treated ZFP143-FKBP cells. **(E)** Genomic tracks showing ZFP143-HA ChIP-seq (red) and nascent transcription (yellow for sense and purple for antisense transcription) measured by TT-seq at *Rpp30* and *Pomgnt2* in DMSO and dTAG-V1 treated ZFP143-FKBP cells. **(F)** Fraction of ZFP143-bound genes among down-regulated (green), up-regulated (pink), and stable (grey) genes in the TT-seq at the indicated times after ZFP143 depletion.

ZFP143-HA peak annotation revealed that most of the detected peaks were located at active promoters^50^ (Figure S1A). *De novo* motif analysis of the peaks revealed two top-scoring motifs, which correspond to the previously reported SBS1 and SBS2 sites^12^ (Figure S1B). These motifs were present in 95% of ZFP143-HA peaks are were among the top enriched TF motifs in the ZFP143-HA peaks^51^ (Figure S1C, S1D), indicating that our ChIP-seq data specifically identifies ZFP143 binding. We compared our ChIP-seq data to a previously published dataset generated using a custom antibody against ZFP143 in J1 mESCs^12^. Our peak set demonstrated a high degree (86%) of overlap with this previously published ZFP143 ChIP-seq mESCs data (Figure S1E). The similarity in the binding profiles of the endogenous and fusion proteins (Figure S1F, Figure S1G), indicates that the introduction of the FKBP12^F36V^ domain and HA tags does not noticeably impact the chromatin occupancy of ZFP143 at its target sites.

To probe the effect of ZFP143 loss on gene transcription, we performed nascent transcript profiling using TT-seq^52^ in control cells and after 2, 6 and 24 hours of ZFP143 depletion. Time course TT-seq analysis revealed that the majority of genes affected by ZFP143 loss are down-regulated (FDR < 0.05, absolute log2-fold-change > 0.5, Figure 1E, S1H), consistent with its role as a transcriptional activator. By combining chromatin occupancy with nascent transcription, we found that 85% of the genes that were down-regulated after 2 hours have ZFP143 binding at the promoter (STAR Methods, Figure 1F), indicating that the down-regulated genes are direct targets of ZFP143. Our findings confirm the functional depletion of ZFP143 and highlight the reliability of our experimental system in detecting the direct effects of ZFP143 loss.

### Acute loss of ZFP143 is not consistent with a role in 3D genome organisation

Recent studies have suggested a role for ZFP143 in 3D genome organisation by contributing to chromatin loop formation^34,38,39,46,47^. To determine the direct consequences of ZFP143 loss on chromatin looping, we performed *in situ* Hi-C following 6 hours of ZFP143 depletion. Analysis of the genome-wide relative contact probability curves showed that they were almost indistinguishable, suggesting no overall changes in genome topology induced by ZFP143 depletion (Figure S2A). To determine whether chromatin loops are affected upon ZFP143 depletion, we performed a pile-up analysis of the loops previously detected in the high-resolution Hi-C mESCs dataset^53^. To our surprise, we did not detect a decrease in genome-wide loop strength (Figure 2A). To investigate whether more specific chromatin loop types might be affected, we analysed cohesin-mediated, enhancer-promoter and promoter-promoter loops^54^. Again, we did not detect a change in contact frequency for any of these loop classes (Figure S2B). Finally, to test whether chromatin loops directly associated with ZFP143 are affected, we analysed subsets of the above loops that had ZFP143 binding at one or both anchors. In this set of loops as well, we were not able to detect any obvious changes in the interaction strength (Figure 2B, S2C).

**Figure 2.**
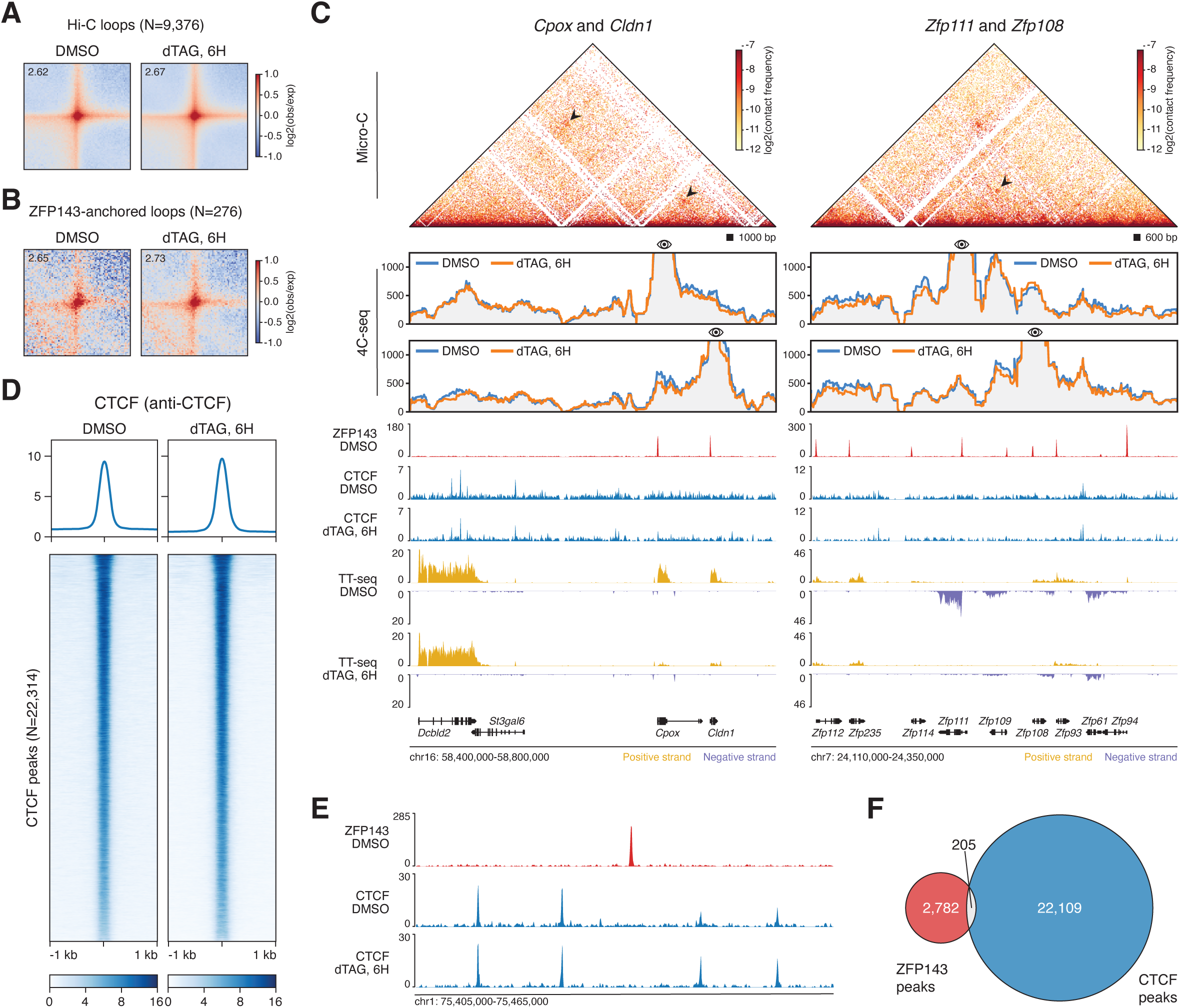
ZFP143 depletion has no detectable effect on 3D genome structure and CTCF binding. **(A)** Average Hi-C loops^53^ in DMSO and 6 hours dTAG-V1 treated ZFP143-FKBP cells. Value in the upper-right corner indicates the interaction strength of the loop over the background. **(B)** Same as in (A) but for the average ZFP143-associated Hi-C loops (containing ZFP143 peak in at least one loop anchor). **(C)** High-resolution 4C-seq data generated for the *Cpox* and *Cldn1* (left panel) and *Zfp111* and *Zfp108* (right panel) loci using gene promoters as viewpoints. The matrix in the top panel represents interaction frequencies in a previously published high-resolution Micro-C dataset^55^. The arrows point to detected Micro-C chromatin loops. The bottom panel shows 4C contact profiles in DMSO (blue) and in 6 hours dTAG-V1 (orange) treated ZFP143-FKBP cells. Genomic tracks show ZFP143-HA ChIP-seq (red), calibrated CTCF ChIP-seq (blue), TT-seq nascent transcription (yellow for sense and purple for antisense transcription) in DMSO and 6 hours dTAG-V1 treated ZFP143-FKBP cells. **(D)** Tornado plots of calibrated CTCF ChIP-seq signal centred at CTCF peaks in DMSO and 6 hours dTAG-V1 treated ZFP143-FKBP cells. **(E)** Genomic tracks showing ZFP143-HA ChIP-seq (red) in DMSO and calibrated CTCF ChIP-seq (blue) in DMSO and 6 hours dTAG-V1 treated ZFP143-FKBP cells. **(F)** Venn diagram showing the overlap between ZFP143-HA (red) and CTCF (blue) peaks.

Puzzled by these findings, we sought to improve the resolution by obtaining the detailed interaction profile of a few loci. To do this, we selected gene promoters that are bound by ZFP143, are downregulated within 2 hours of ZFP143 depletion, and form a loop in a recently published high-resolution Micro-C mESCs dataset^55^. We performed 4C-seq^56^ after 6 hours of ZFP143 depletion, using the selected promoters as viewpoints. Similar to the genome-wide Hi-C results, the 4C-seq interaction profiles of these regions were mostly unaffected by ZFP143 depletion for any of the promoters tested (Figure 2C, S2D). Although some subtle changes in the interaction frequencies were observed for several viewpoints, it is more likely that these are a consequence of the transcriptional changes rather than a direct effect of ZFP143 depletion. Overall, our comprehensive analyses do not detect systematic changes in the chromatin interaction landscape of mESCs following ZFP143 depletion. Therefore, our results are not consistent with a role for ZFP143 in establishing or maintaining genome architecture.

### CTCF binding and distribution is unaffected upon ZFP143 depletion

CTCF is a key factor in mediating chromatin contacts by creating a barrier to cohesin extrusion^29^. Recently, a knock-out of ZFP143 in mouse haematopoietic stem and progenitor cells suggested a loss of CTCF binding in the regions where ZFP143 and CTCF colocalise^38^, which would implicate ZFP143 in positioning CTCF on chromatin. To determine whether we could detect a similar dependency following acute depletion of ZFP143 in mESCs, we performed a calibrated CTCF ChIP-seq after 6 hours of ZFP143 loss. Surprisingly, we could not detect any consistent change in CTCF occupancy after 6 hours of ZFP143 depletion (Figure 2D). Heatmaps of ZFP143 binding intensity at CTCF peaks demonstrated hardly any ZFP143 signal at these regions (Figure S2E). Moreover, very little CTCF signal was detected at the ZFP143 bound sites (Figure S2F). Visual inspection of the chromatin occupancy tracks indicated that CTCF and ZFP143 binding were largely mutually exclusive (Figure 2E), and the calculated overlap between CTCF and ZFP143 peaks showed that only a minority of them were shared (Figure 2F). In contrast to previous reports, these results suggest that depletion of ZFP143 does not affect CTCF positioning and that ZFP143 and CTCF bind to distinct genomic regions.

### Antibody cross-reactivity led to discovery of ZNF143 colocalization with CTCF and subsequent role in chromatin looping

The results we obtained from our acute depletion of ZFP143 appear to be at odds with the literature reports. To determine the source and nature of the discrepancies, we systematically re-analysed published ZNF143/ZFP143 and CTCF ChIP-seq data. We decided to overlap ZNF143/ZFP143 peaks from publicly available datasets with CTCF peak sets available in the CISTROME database^57^ (Table S2). For each dataset, we calculated the proportion of ZNF143/ZFP143 peaks that overlapped with CTCF peaks, regardless of cell type (STAR Methods, Figure 3A, Table S3). For the majority of ZNF143/ZFP143 datasets, the median proportion of peaks overlapping with CTCF peaks was approximately 20%. However, we found that for certain datasets the percentage of overlap is much higher (>40%) (Figure 3A). The datasets that showed the greatest overlap with CTCF peaks were obtained using the same antibody, anti-ZNF143 Proteintech 16618-1-AP (hereafter referred to as Proteintech), which is a polyclonal antibody that recognises the endogenous ZNF143 protein. In addition, most of these datasets originate from the ENCODE project, in which the overlap with CTCF was first reported^30^.

**Figure 3.**
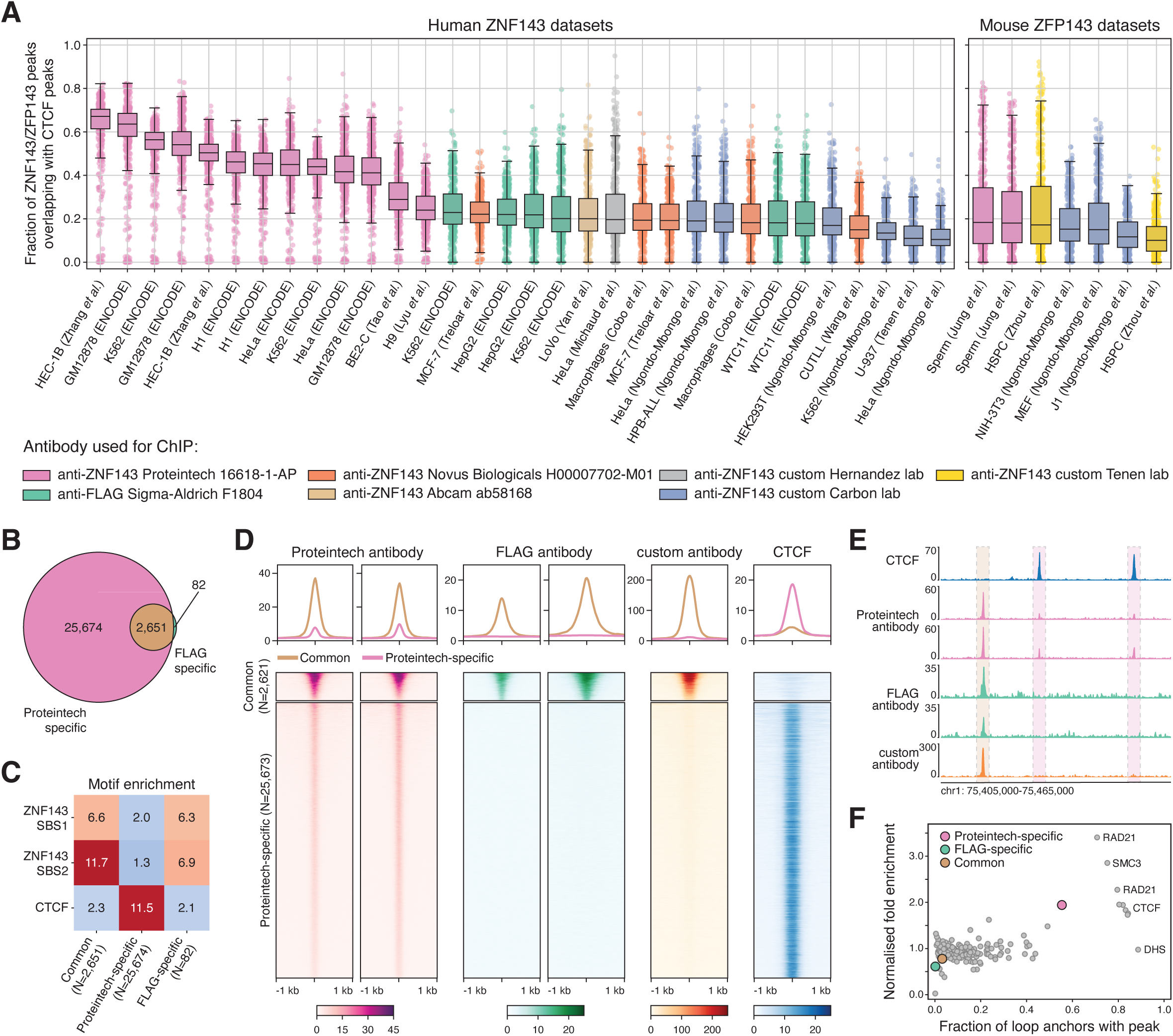
Re-analysis of publicly available ChIP-seq data reveals ZNF143 antibody cross-reactivity with CTCF. **(A)** Overlap between ZNF143/ZFP143 peaks from re-analysed publicly available data and CTCF peaks from CISTROME^57^ for human (left panel) and mouse (right panel) datasets. Box plots for each ZNF143/ZFP143 dataset represent the median overlap with CTCF peaks. Each dot represents the overlap between the indicated ZNF143/ZFP143 peak set with an individual CTCF peak set. Colours represent the antibody used for chromatin immunoprecipitation, as indicated below. **(B)** Venn diagram showing the overlap between ZNF143 peaks detected by Proteintech (light pink) and FLAG (light green) antibodies in K562 cells. **(C)** Heatmap showing the enrichment of SBS (i.e. ZNF143) and CTCF motifs in common, Proteintech-specific, and FLAG-specific peaks in K562 cells. (D) Tornado plots of ChIP-seq signals detected by Proteintech (light pink), FLAG (light green), and custom^12^ (orange) antibodies, and CTCF signal (blue) in K562 cells. The ChIP-seq signals are centred on common (top) and Proteintech-specific (bottom) peaks. (E) Genomic tracks showing ChIP-seq signals for CTCF (blue) and signals detected by Proteintech (pink), FLAG (light green), and custom12 (orange) antibodies in K562 cells. Rectangles indicate common (left) and Proteintech-specific (middle and right) peaks in the region. (F) Scatter plot of the percentage of loop anchors overlapping the peak (x-axis) against the fold enrichment of peaks in loop anchors (y-axis) for a number of DNA binding proteins^40^ and for Proteintech-specific, FLAG-specific and common peaks in K562 cells.

To further explore the overlap between ZNF143 and CTCF, we compared ChIP-seq data generated using Proteintech antibody (anti-ZNF143) with C-terminally 3xFLAG-tagged ZNF143 ChIP-seq (anti-FLAG-ZNF143) in K562 cell line^58^. Surprisingly, we found that there was almost a 10-fold difference in the number of peaks picked up by the Proteintech and FLAG antibodies. Despite this difference, the anti-FLAG-ZNF143 peaks are an almost complete subset of the anti-ZNF143 peaks (Figure 3B). However, there were 25,673 peaks identified only with the Proteintech antibody (Proteintech-specific), in contrast to 2,621 regions recognised by both FLAG and Proteintech (common) and 82 picked up only by the FLAG antibody (FLAG-specific) (Figure 3B). We used these three peak categories to perform a motif enrichment analysis with the identified ZNF143 SBS1 and SBS2 motifs and the CTCF motif from the JASPAR database^59^. The two ZNF143 SBS motifs were strongly enriched in the common and FLAG-specific categories, but showed low enrichment in the Proteintech-specific category of peaks (Figure 3C). On the other hand, the CTCF motif was highly enriched in the Proteintech-specific category and only slightly enriched in the other two categories (Figure 3C). Visual inspection and genome-wide quantification of the chromatin occupancy tracks showed that the Proteintech anti-ZNF143 antibody detected signal at CTCF binding sites, whereas nothing could be detected in the anti-FLAG-ZNF143 data at the same regions (Figure 3D, 3E). Additionally, we included the data generated with a custom anti-ZNF143 antibody from a separate study^12^, which showed a similar binding profile to anti-FLAG-ZNF143, with strong binding at the common peaks and lack of binding at the CTCF-bound Proteintech-specific peaks (Figure 3D, 3E). It should be noted that the signal detected with the Proteintech antibody at the Proteintech-specific peaks has a much lower intensity compared to its signal at the common peaks (Figure 3D, 3E). These results strongly suggest that the data obtained with the anti-ZNF143 Proteintech antibody has an internal bias, which may be explained by antibody-cross reactivity with CTCF.

CTCF is strongly associated with chromatin looping and loss of CTCF results in loss of CTCF-anchored cohesin-mediated chromatin loops^60^. Systematic analysis of 3D genome organisation using *in situ* Hi-C revealed that ZNF143, together with CTCF and cohesin subunits, is highly enriched at chromatin loop anchors in GM12878 and K562 cell lines^40^. We decided to recalculate the enrichments of proteins at the loop anchors defined in the K562 Hi-C data using the three ZNF143 peak categories mentioned above. We found that only the Proteintech-specific peaks show a high enrichment at loop anchors, while the other two categories do not (Figure 3F). Therefore, we suspect that the peaks detected by the Proteintech antibody, which do not show enrichment of the ZNF143 motif and are strongly associated with CTCF binding, explain the reported enrichment of ZNF143 at chromatin loop anchors. These analyses challenge the idea that ZNF143 has a role in chromatin looping.

Finally, there have been several reports of chromatin binding interdependence between ZNF143 and CTCF, namely two opposing claims that CTCF is positioned on chromatin by ZNF143^38^ and that ZNF143 occupancy depends on CTCF at regions where the two proteins bind together^39^. Re-analysis of the data from these studies suggests that the reduction of CTCF on chromatin after ZNF143 knockout may have been caused by GC bias in the ChIP-seq data (STAR Methods, Figure S3A-F), whereas the reduction of ZNF143 occupancy following CTCF depletion may be explained by the use of the Proteintech antibody that recognises CTCF due to cross-reactivity (STAR Methods, Figure S3G-H). In addition, we found that all ZNF143-CTCF motif pairs, reported to localise in 37 bp proximity to each other^38^, are contained within SINE/B2 repetitive elements and are likely to be an artefact (STAR Methods, Figure S3I-J). Taken together, these results suggest that cooperation between ZNF143 and CTCF is highly unlikely.

### ZFP143 regulates nuclear genes encoding mitochondrial proteins

Our results so far suggest that ZFP143 is not involved in chromatin looping, and we therefore focused on understanding its role as a transcription factor. Since ZFP143 binds predominantly to transcription start sites, we used the ZFP143-HA ChIP-seq data to identify ZFP143-bound gene promoters in mESCs (N=2226). Gene ontology analysis of ZFP143-bound genes revealed that they belong to functional categories previously associated with ZFP143^12^, such as snRNAs, cell cycle, and metabolism (Figure S4A, Table S4). In addition, we found a number of gene sets involved in translation, mitochondrial functions and stress-related pathways (Figure S4A). To understand the conservation of ZNF143/ZFP143-bound genes, we utilised the systematically re-analysed ZNF143/ZFP143 ChIP-seq data. We identified a set of ZNF143/ZFP143 binding sites conserved across human and mouse cells (STAR Methods, Figure S4B) and associated them to gene promoters (1696 and 1653 conserved ZNF143/ZFP143-bound genes in human and mouse, respectively, STAR Methods). Overlapping the identified conserved ZNF143/ZFP143-bound genes in human and mouse with the ZFP143-bound genes from our study revealed a strong overlap between the genes (54% overlap conserved human ZNF143-bound genes, 67% overlap conserved mouse ZFP143-bound genes, Figure S4C) and the functional categories to which they belong (Figure S4D, S4E, Table S4). This suggests that regulation by ZNF143/ZFP143 is highly conserved across cell types and organisms.

To analyse the direct functional consequences of ZFP143 loss, we performed a more detailed analysis of our time course TT-seq dataset following ZFP143 depletion. Differential expression analysis identified several hundred genes for which transcription was down-regulated (FDR < 0.05, absolute log2-fold-change > 0.5, Figure 4A, Table S5), and the number of down-regulated genes increased with longer depletion times. In the first 6 hours following ZFP143 depletion only a handful of genes increased their expression, and only after 24 hours did we observe hundreds of up-regulated genes (Figure 4A). The lack of ZFP143 binding to the promoters of these genes (Figure 1F) and the late response indicates that the effect of ZFP143 depletion on the expression of these genes is indirect. Of note, not all ZFP143-bound genes are down-regulated, suggesting that ZFP143 directly and exclusively controls only a subset of these, while the others may depend on additional factors for their activation.

**Figure 4.**
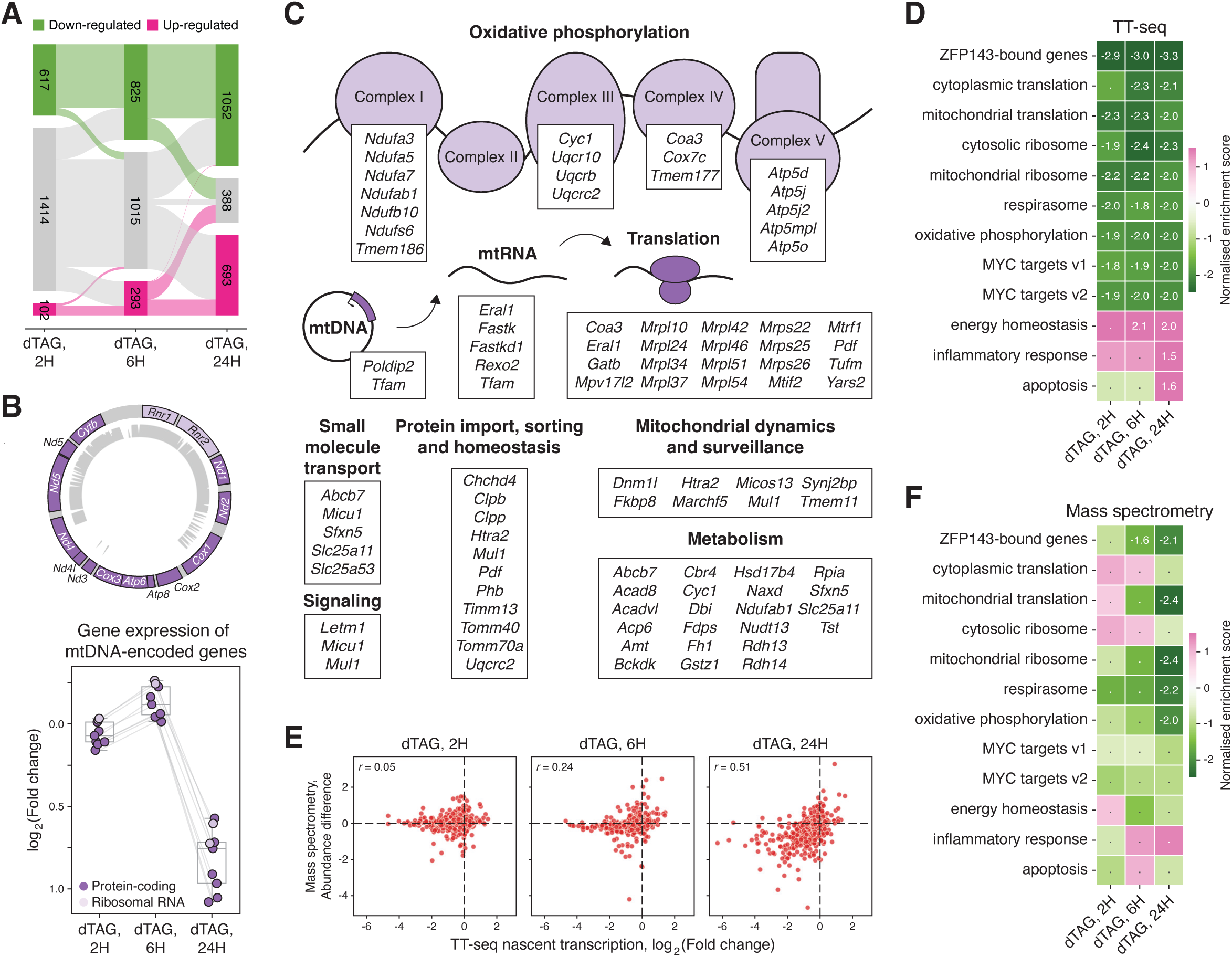
ZFP143 depletion leads to down-regulation of mitochondria-related genes at transcriptome and proteome levels. **(A)** Sankey diagram showing the differentially expressed genes following the time course of ZFP143 depletion for 2, 6 and 24 hours: up-regulated (pink) and down-regulated (green) genes at FDR < 0.05, absolute log2-fold-change > 0.5. **(B)** Schematic of the genes and single-read mappability^116^ (k=36) for mitochondrial DNA (top panel) and fold changes of the mitochondrial-encoded genes detected by TT-seq following ZFP143 depletion time course (bottom panel). **(C)** Schematic of nuclear-encoded mitochondrial genes that are bound by ZFP143 in ChIP-seq data and are down-regulated in TT-seq data. Genes are grouped by mitochondrial function or protein complex as defined in the MitoCarta database^62^. **(D)** Gene set enrichment analysis of TT-seq data after ZFP143 depletion (upregulated gene sets in pink, down-regulated gene sets in green). The normalised enrichment score for significant gene sets (FDR < 0.05) is shown. **(E)** Correlation of nascent RNA production measured by TT-seq (x-axis) and protein abundance measured by mass spectrometry (y-axis) for ZFP143-bound genes in 2, 6, and 24 hours dTAG-V1 treated ZFP143-FKBP cells. Values in the upper-left corners indicate Pearson correlation coefficients. **(F)** Gene set enrichment analysis of mass spectrometry data after ZFP143 depletion (upregulated gene sets in pink, down-regulated gene sets in green). The normalised enrichment score for significant gene sets (FDR < 0.05) is shown.

Inspection of differentially transcribed genes revealed that all protein-coding and rRNA genes from the mitochondrial genome detected by TT-seq show significant down-regulation after 24 hours of ZFP143 loss (Figure 4B). Consistent with this finding, *Tfam*, which encodes an essential mitochondrial transcription factor^61^, is down-regulated after 2 hours of dTAG-V1 treatment and is a target of ZFP143. To investigate whether other nuclear-encoded genes involved in mitochondria-related processes are specifically down-regulated upon ZFP143 loss, we annotated differentially expressed genes using the MitoCarta database^62^ and found multiple examples that show similar behaviour. These include subunits of multiple oxidative phosphorylation complexes *Ndufab1*, *Cyc1*, and *Coa3*, translation factors *Mtrf1* and *Tufm*, mitochondrial fission regulator *Dnm1l*, components of the protein import TIM/TOM complex, NAD metabolism genes, and others^61,63^ (Figure 4C, Table S6). To gain further insight into which gene categories are deregulated upon ZFP143 depletion, we performed gene set enrichment analysis (GSEA) on the TT-seq data, which revealed that gene categories involved in cytoplasmic and mitochondrial translation, oxidative phosphorylation components and MYC targets show significant down-regulation (Figure 4D, Table S7). These gene sets were also identified as significantly enriched for ZFP143 binding genes in our analyses (Figure S4A, S4E). We found hardly any gene sets showing significant up-regulation at the early time points, although after 24 hours of ZFP143 depletion, differentiation-related gene sets, as well as apoptosis and inflammatory response categories show significantly increased expression (Figure 4D, Table S7), suggesting activation of a general stress response in mESCs.

To investigate how transcriptional changes translate into changes at the protein level, we performed quantitative mass spectrometry-based proteome analysis. We sampled the same time points following ZFP143 depletion as in the TT-seq data. As a control, we first treated untagged parental mESCs with dTAG-V1 and identified only minimal changes in the proteome, indicating that dTAG-V1 treatment does not appreciably affect protein abundance (Figure S5A). In ZFP143-depleted mESCs, the composition of the proteome is largely unaffected at 2 and 6 hours after depletion. However, at 24 hours after ZFP143 depletion, the levels of hundreds of proteins are significantly reduced (Student’s t-test –log(p-value) ≥ 1.3, absolute abundance difference ≥ 0.5, Figure S5B, Table S8), 73% of which are encoded by genes bound by ZFP143 at the promoter (Figure S5C), suggesting a direct relationship between gene regulation by ZFP143 and the abundance of gene products in the proteome. To assess the temporal relationship between transcription and protein levels, we compared the changes in nascent RNA production measured by TT-seq and protein abundance measured by mass spectrometry. For ZFP143-bound genes, the decrease in nascent transcription shows strong correlation (Pearson r=0.51, p-value<2.2e-16) with a decrease in protein abundance only after 24 hours of ZFP143 loss (Figure 4E). Our results indicate that ZFP143-bound genes that are down-regulated at early time points are manifested in the proteome at later time points, consistent with delayed functional consequences.

To determine which proteins are affected by ZFP143 depletion, we performed GSEA and compared it with the results obtained from TT-seq (Figure 4F, Table S9). At 2 hours post depletion not a single gene set demonstrates a significant change in protein abundance. At 6 hours post depletion, ZFP143-bound genes are the only category of proteins that shows a significant decrease in abundance. Additionally, at 24 hours, besides ZFP143-bound genes, mitochondrial translation and oxidative phosphorylation functional categories also become significantly down-regulated in the proteome, consistent with the TT-seq outcome. These results show that ZFP143 is a transcriptional regulator of nuclear-encoded mitochondrial genes and that its depletion has an immediate effect on their transcription and a direct downstream effect at the protein level.

### Prolonged ZFP143 loss leads to decreased cell proliferation and mitochondrial dysfunction

To probe the consequences of ZFP143 loss on cell morphology and proliferation, we monitored mESCs over a longer period by extending dTAG-V1 treatment to 24, 48 and 72 hours. We found that after 24 hours of ZFP143 loss the morphology of the mESCs colonies changes profoundly (Figure 5A). The colonies lose their dome-like structure and tend to become more flattened out, and this effect becomes particularly clear after 72 hours of dTAG-V1 treatment. In addition, cells depleted of ZFP143 showed a clear reduction in cell number compared to non-depleted DMSO-treated cells (Figure 5B). We hypothesised that the reduced cell number could be a consequence of (i) cell cycle arrest, (ii) slower proliferation rate or (iii) increased cell death. To determine whether there was an effect on the cell cycle we performed cell cycle analysis using a 5-ethynyl-2′-deoxyuridine (EdU) incorporation assay. We did not observe a shift in the cell cycle phase following ZFP143 depletion (Figure S6A, S6B), meaning that cells do not arrest at a specific point in the cell cycle. However, we did observe a decrease in the EdU signal intensity for the S phase cells, suggesting lower EdU incorporation rates (Figure S6C). This observation is consistent with ZFP143-depleted cells spending more time on DNA replication and by extension a slower proliferation rate. To determine whether there was increased cell death through apoptosis, we performed Annexin V staining on the control and dTAG-V1 treated cells (Figure S6D). The percentage of Annexin V-positive cells was slightly elevated at 72 hours following ZFP143 depletion, but not at the earlier time points (Figure S6E, S6F). These findings imply that ZFP143 is essential for maintaining a high proliferation rate of mESCs.

**Figure 5.**
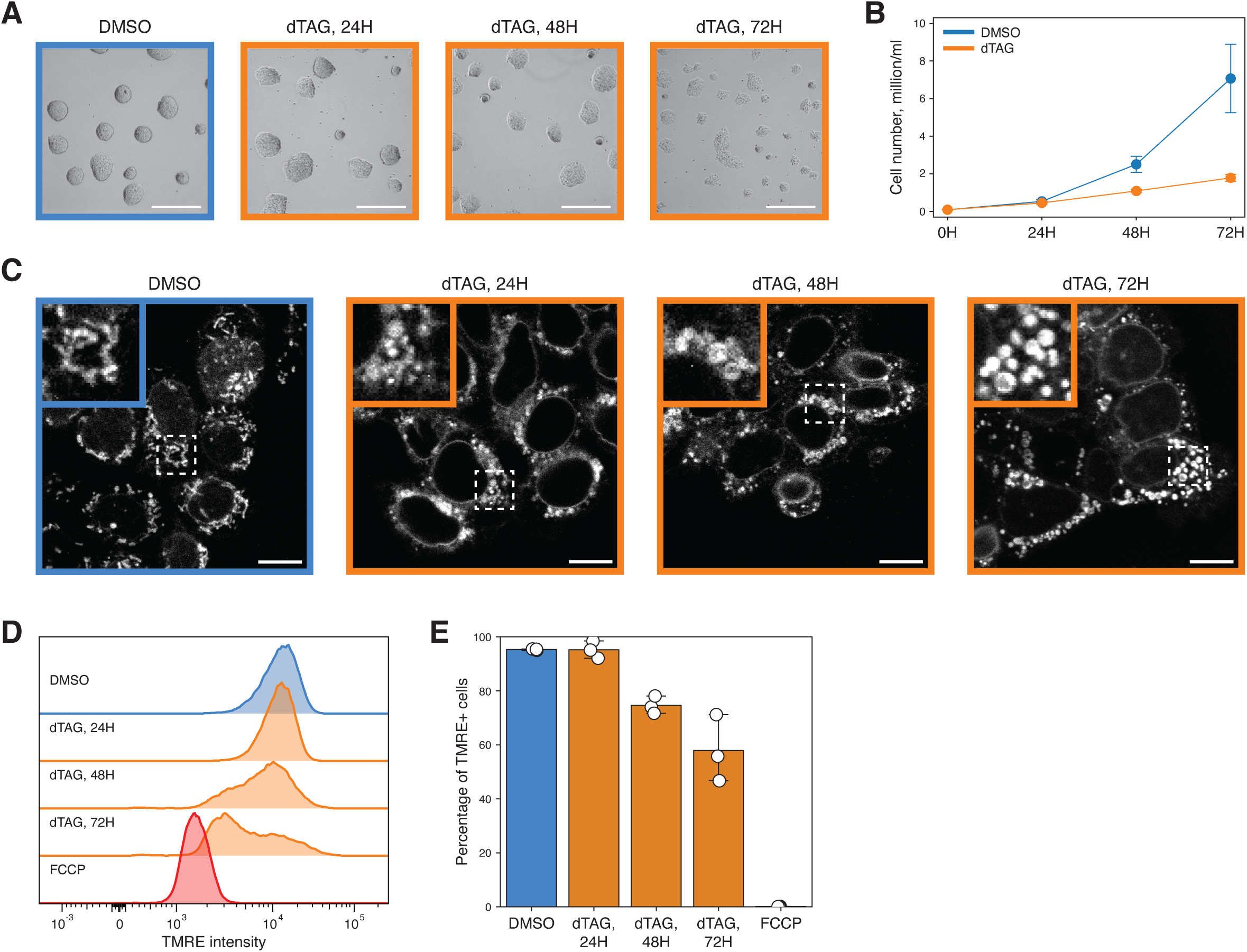
Prolonged loss of ZFP143 impairs cell proliferation and mitochondrial function. **(A)** Bright-field microscopy images showing morphology of DMSO (blue frame) and 24, 48, and 72 hours dTAG-V1 treated (orange frames) ZFP143-FKBP mESC colonies. Scale bar: 200 µm. **(B)** Growth curve showing the total cell number of DMSO (blue line) and dTAG-V1treated (orange line) ZFP143-FKBP cells at different time points of treatment. Dots indicate mean values. Error bars indicate standard deviation. **(C)** Confocal microscopy images of the MitoTracker Red FM fluorescence in DMSO (blue frame) and 24, 48, and 72 hours dTAG-V1 treated (orange frames) ZFP143-FKBP cells. Magnified images of mitochondrial morphology and mitochondrial network for the dash-boxed regions in the representative images are shown in the upper-left corners. Scale bar: 10 µm. **(D)** Representative flow cytometry histogram showing TMRE fluorescence distribution in DMSO (blue), 24, 48, and 72 hours dTAG-V1 (orange), and FCCP (red) treated ZFP143-FKBP cells. **(E)** Quantification of TMRE-positive cells in DMSO (blue), 24, 48 and 72 hours dTAG-V1 (orange), and FCCP (red) treated ZFP143-FKBP cells. Dots represent values for replicates. Error bars indicate 95% confidence interval.

Our transcriptomic and proteomic analyses showed that ZFP143 maintains expression of nuclear-encoded mitochondrial genes. Since mitochondria are involved in the regulation of proliferation, cell cycle and cell death^64,65^, we decided to address the effect of ZFP143 loss on mitochondrial function. To this end, we visualised mitochondria in living cells using the fluorescent MitoTracker probe^66^. In control mESCs we observed elongated mitochondria and mitochondrial networks with high connectivity (Figure 5C). Strikingly, ZFP143-depleted cells present increased amounts of larger, circular mitochondria that have fewer interactions with each other (Figure 5C). Such morphological changes have been associated with changes in mitochondrial activity^67–69^, so we decided to measure the mitochondrial membrane potential. For this, we utilised positively charged fluorescent tetramethylrhodamine ethyl ester (TMRE) dye, which accumulates in polarised mitochondria and can be used to assess their function^70^. Flow cytometry measurements of the TMRE signal demonstrated that ZFP143 depletion resulted in a gradual loss of TMRE intensity (Figure 5D, S6G). The TMRE signal was shifted towards the positive control obtained by treating cells with the mitochondrial uncoupler FCCP^71^, indicating that more depolarised mitochondria were present (Figure 5D, S6G). In terms of temporal dynamics of mitochondrial membrane potential loss, we did not observe any changes in the mitochondrial membrane potential of the cells at 24 h post-depletion, despite the alterations in the mitochondrial proteome and mitochondrial morphology. However, after 48 hours of dTAG-V1 treatment, we observed a drop in the percentage of TMRE-positive cells, which further decreased at 72 hours post depletion, corresponding to an increase of cells with depolarised mitochondria (Figure 5E). Notably, none of the mitochondrial phenotypes described above were observed for the untagged parental mESCs treated with dTAG-V1 (Figure S7). Taken together, the disrupted cell proliferation, altered mitochondrial morphology and loss of mitochondrial membrane potential collectively underline the stress conditions induced by long-term ZFP143 depletion.

### ZFP143-depleted cells fail to form stem cell-derived embryo models

Studies in zebrafish embryogenesis and mouse haematopoiesis have implicated ZNF143 in development^24,26,38^. However, why ZNF143 is essential for organismal development has remained elusive. To investigate its role in mammalian development, we used an *in vitro* embryo model, called gastruloids, which mimics early stages of post-occipital embryonic development^72,73^. Gastruloids are generated by aggregating embryonic stem cells and providing a pulse of the Wnt agonist Chiron, which promotes symmetry breaking and formation of multiple cell types. We grew gastruloids from the ZFP143-FKBP mESC line and depleted ZFP143 by treating them with dTAG-V1 at different times during gastruloid development. Strikingly, ZFP143 depletion at 48 hours post aggregation dramatically impaired gastruloid formation (Figure 6A). Compared to DMSO-treated control gastruloids, ZFP143-depleted gastruloids failed to elongate and produced a small spherical aggregate at 120 hours (Figure 6A, 6B). Treatment with dTAG-V1 at 72 hours of gastruloid formation produced a milder phenotype, although these gastruloids were smaller and less elongated compared to control gastruloids. Treatment at 96 hours produced gastruloids morphologically indistinguishable from the control (Figure 6A, 6B).

**Figure 6.**
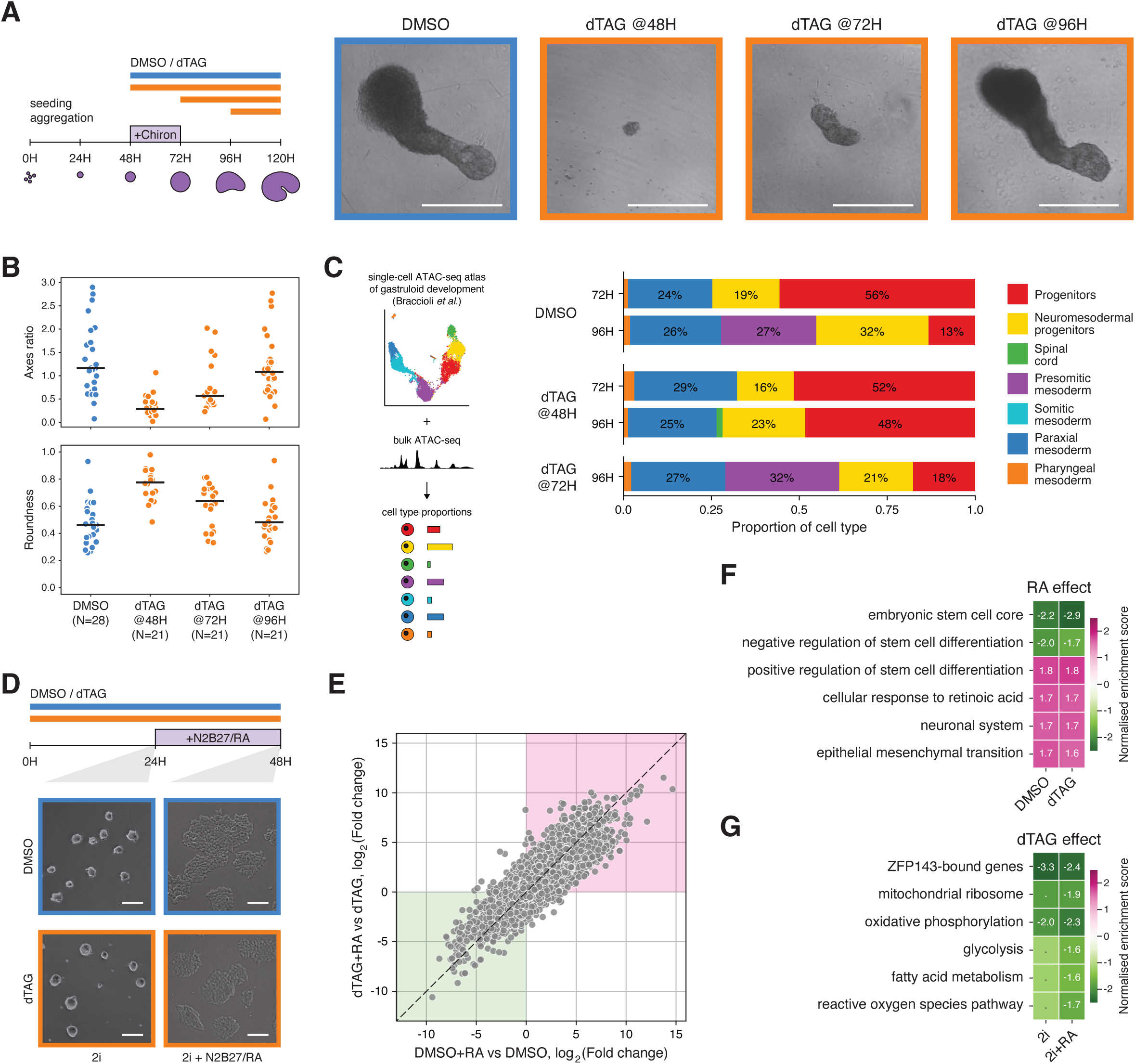
ZFP143-depleted cells fail to develop fully differentiated embryo models despite being able to exit the pluripotent state. **(A)** Schematic of the *in vitro* gastruloids experiment (left panel). Bright-field microscopy images showing gastruloids treated with DMSO (blue) and dTAG-V1 from 48, 72, and 96 hours stages (orange), imaged at 120h of development (right panel). Scale bar: 300 µm. **(B)** Quantification of axis ratio (top panel) and roundness (bottom panel) of gastruloids treated with DMSO (blue) and dTAG-V1 from 48, 72, and 96 hours stages (orange), imaged at 120h of development. Black lines indicate the mean values. The number of quantified gastruloids is indicated below. **(C)** Schematic of the ATAC-seq deconvolution pipeline (left panel). Quantification of cell type proportions using ATAC-seq deconvolution in gastruloids treated with DMSO or dTAG-V1 at 48 and 72 hours stages (right panel). ATAC-seq was performed at 72 and 96 hours stages. Colours represent cell-type annotation as referred to in the original study^74^. **(D)** Schematic of the retinoic acid differentiation experiment (top panel). Bright-field microscopy images showing morphology of DMSO (blue frames) and dTAG-V1 treated (orange frames) ZFP143-FKBP mESC colonies grown for 24 hours in 2i media before (left) and after (right) transfer to N2B27 differentiation media with retinoic acid for 24 more hours (bottom panel). Scale bar: 100 µm. **(E)** Correlation of gene expression changes measured by RNA-seq in DMSO and dTAG-V1 treated ZFP143-FKBP cells differentiated with retinoic acid. Each dot represents a single gene, pink and green areas represent concordantly up- and down-regulated genes, respectively. **(F)** Gene set enrichment analysis of RNA-seq data in ZFP143-FKBP mESCs, showing the effect of retinoic acid in DMSO and dTAG-V1 treated cells (up-regulated gene sets in pink, down-regulated gene sets in green). The normalised enrichment score for significant gene sets (FDR < 0.1) is shown. **(G)** Gene set enrichment analysis of RNA-seq data in ZFP143-FKBP mESCs, showing the effect of dTAG-V1 treatment before (2i) and after (2i+RA) transfer to differentiation media (up-regulated gene sets in pink, down-regulated gene sets in green). The normalised enrichment score for significant gene sets (FDR < 0.1) is shown.

To understand the growth dynamics of gastruloids in the presence and absence of ZFP143, we performed time-course imaging at the intermediate time points. We found that gastruloids depleted of ZFP143 at 48 hours can develop normally up to the 72 hours time point, after which they substantially reduce in size and ultimately collapse (Figure S8A, S8B, see 96 hours imaging). Similarly, gastruloids treated with dTAG-V1 at the 72 hours time point grow at a similar pace as control gastruloids up to 96 hours after which they shrink (Figure S8A, S8B). These observations show that in the absence of ZFP143, gastruloids continue to grow normally for 24 hours, after which they begin to shrink and even collapse.

We next sought to determine the effect of ZFP143 loss on the cell type composition of gastruloids. We focused on gastruloids depleted of ZFP143 at 48 and 72 hours of development and performed bulk chromatin accessibility measurements by ATAC-seq. To estimate cell type proportions, we deconvolved the bulk ATAC-seq signal using a time-resolved single-cell chromatin accessibility atlas of gastruloid development that we previously generated^74^. Deconvolution analysis shows that at 72 hours of development, control DMSO-treated gastruloids are mainly composed of progenitor cells, resembling epiblast-like state with regulatory landscape dominated by the activity of OCT4 and SOX2^74,75^ (Figure 6C). This progenitor population is dramatically reduced by 96 hours, suggesting that the cells are exiting the pluripotent state and give origin to more differentiated cell types (Figure 6C). However, in gastruloids depleted of ZFP143 at 48 hours of development, progenitor cells still make up the largest cell population at 96 hours, whereas in gastruloids depleted of ZFP143 at 72 hours we observed the same cell type proportions as in control ones (Figure 6C). These results suggest that ZFP143 may be required for the differentiation of pluripotency-associated progenitors in gastruloid development.

We then sought to understand whether ZFP143 is required to maintain the differentiation capacity of stem cells by facilitating the exit from the pluripotent state. To this end, we asked whether ZFP143-depleted mESCs could differentiate in 2D culture upon treatment with retinoic acid (RA). RA is widely used to differentiate mESCs into neuronal lineages *in vitro*^76^ and plays an important role in embryonic development^77^. We first induced ZFP143 degradation by pre-treating mESCs with either dTAG-V1 or DMSO for 24 hours, and then induced differentiation by adding RA for the next 24 hours while keeping the cells in dTAG-V1 or DMSO, respectively. Treatment of cells with RA had no further effect on the proliferation rates of both control and ZFP143-depleted mESCs (Figure S9A), and they both were able to change their morphology into more elongated neural-like cells, indicating successful RA-induced neuronal differentiation (Figure 6D).

To determine whether differentiation with RA induces similar transcriptomic changes, we performed RNA-seq before and after RA treatment and compared the molecular effects on differentiation in control and ZFP143-depleted cells. Principal component analysis showed that RA treatment induces a similar trend of changes in control and dTAG-V1 treated mESCs, accounting for 83% of the variance between samples, while the difference induced by ZFP143 depletion represents only 14% of the variance (Figure S9B). To formally compare RA-treated control and ZFP143-depleted cells, we performed differential expression analysis (Figure S9C, Table S10), which revealed that differentially expressed genes in both conditions have a high degree of overlap and show concordant changes in expression (Figure 6E, S9D). Furthermore, GSEA showed that both control and ZFP143-depleted cells show a reduced expression of stem cell-related genes and an upregulation of genes involved in RA response, neural lineage differentiation and epithelial-to-mesenchymal transition (Figure 6F, Table S11), demonstrating that the changes in transcriptional programmes with and without ZFP143 upon RA stimulation are highly similar. However, examining the effects of ZFP143 depletion in cells before and after RA treatment, revealed down-regulation of nuclear-encoded mitochondrial genes involved in energy metabolism and redox homeostasis (Figure 6G, Table S11). This confirms that ZFP143 is not directly required for stem cell differentiation in monoculture, but maintains a cellular state that drives proper organismal development in a multicellular context.

## Discussion

In this study, we investigated the molecular consequences of acute loss of ZFP143. To our surprise, we were unable to identify a direct role for ZFP143 in chromatin looping, despite numerous studies suggesting that this protein is a looping factor. Our detailed analyses of published ChIP-seq and Hi-C data show that the role of ZFP143 as a looping factor is most likely a consequence of cross-reactivity of a polyclonal anti-ZNF143 antibody with the *bona fide* chromatin looping factor CTCF. By profiling the nascent transcriptome, we found that ZFP143 is involved in the direct transcriptional activation of hundreds of genes. Our results show that ZFP143 binds to promoters of a conserved set of genes involved in various mitochondrial processes, such as oxidative phosphorylation, mitochondrial translation, and others. Depletion of ZFP143 leads to immediate down-regulation of its target genes, suggesting that ZFP143 acts as a direct transcriptional regulator of nuclear-encoded mitochondrial genes. Prolonged loss of their expression results in mitochondrial dysfunction, decreased proliferation and developmental defects.

We were unable to identify a role for ZFP143 in chromatin looping by examining the genome-wide interaction landscape using Hi-C and the interactions of a selected set of ZFP143 binding sites using 4C-seq following acute loss of ZFP143. An accompanying study (<u>Narducci and Hansen</u>) found that even in nucleosome-resolution chromosome conformation maps generated with Micro-C, no consistent changes in chromatin looping could be observed in either mESCs or human HEK293T cells upon acute ZFP143 loss. It should be noted that previous studies suggesting a role for ZNF143 in chromatin looping show only mild effect sizes, particularly when compared to depletion of well-established regulators of chromatin organisation such as CTCF^60,78^, cohesin^79,80^, WAPL^81,82^ or NIPBL^83^. Furthermore, the reported changes were detected in a non-acute ZNF143 depletion setting^38,39,46,47^, which means that any changes that were observed could also be an indirect consequence of prolonged ZNF143 loss. After systematically re-analysing the ZNF143 ChIP-seq data, we posit that the Proteintech anti-ZNF143 polyclonal antibody recognises CTCF in addition to ZNF143. To our knowledge, the only studies that report an overlap between ZNF143 and CTCF are the ones that utilised this antibody^30,31,33–35^. The use of the same datasets led to the observed enrichment of ZNF143 at chromatin loop anchors^35,36,40^ and thus its misassociation with chromatin looping. Our findings question the use of polyclonal antibodies for ChIP-seq experiments^84^ and emphasise the need for vigilance when interpreting the data generated with polyclonal antibodies, particularly for chromatin factors.

By combining time-resolved transcriptional changes induced by acute ZFP143 loss with ZFP143 chromatin binding, predominantly at gene promoters, we were able to confidently identify direct targets of ZFP143 in our degron mESC line. We found that loss of ZFP143 leads to down-regulation of hundreds of genes, the majority of which have ZFP143 binding at the transcription start site. Moreover, we observed a profound, yet delayed, correlation between transcription of these genes and their protein abundance levels. As in general there is a lack of concordance between transcriptome and proteome^85^, a tight relation between the two upon ZFP143 degradation is remarkable and may indicate a global housekeeping function for ZFP143^86–89^. As for the functional annotation of ZFP143 target genes, we found that many of them are nuclear-encoded mitochondrial proteins, including components of mitochondrial ribosome and electron transport chain complexes. While post-transcriptional regulation of mitochondrial proteins and mitochondrial structure and dynamics is relatively well studied^90–92^, little is known about the transcriptional regulation of these genes in the nucleus, and our dataset can serve as a useful starting point for future research on this topic.

From our analyses, we found that most ZFP143-bound genes show no change in their expression following depletion, indicating that ZFP143 is not the primary activator at many binding sites. The role of ZFP143 at these bound but not regulated sites is an interesting question for future research. Given that these sites partially comprise a conserved set of ZNF143/ZFP143-bound promoters between cell types and even species, it seems likely that these regions are functional at the organismal level. One possibility is that ZFP143 is required for activation at some of these sites in different cell types thus posing its role as a cell-type-specific transcription factor. Another explanation may be that TFs, such as HCFC1, THAP11 and others, which overlap with ZFP143 binding^12,19,93–95^, cooperate or compete with ZFP143 at these promoters and may take over the regulatory function in its absence. Additionally, ZFP143 binding sites were also found to overlap with ICN1, the active form of the transmembrane receptor NOTCH1^12,96^, suggesting it may be involved in signalling responses. Together with the auto-regulatory mechanism of ZFP143 transcriptional levels^97^ and long chromatin residence time at its target sites (<u>Narducci and</u> <u>Hansen</u>), this poses an interesting hypothesis that ZFP143 may be required for rapid transcriptional responses to accommodate specific metabolic conditions and developmental cues.

We have also investigated the effects of prolonged ZFP143 depletion on mESCs. We found that depletion of ZFP143 for more than 24 hours resulted in reduced proliferation and mitochondrial dysfunction. The observed phenotype is similar to the direct effects on mitochondrial function following inhibition of mitochondrial translation^98–100^, blocking the functionality or proper assembly of oxidative phosphorylation complexes^98,100–102^, or shifting the balance between nuclear and mitochondrial-encoded mitochondrial proteins^103,104^. Additionally, we observed reduced proliferation, consistent with a prolonged S phase in our degron line. ZNF143 loss has previously been associated with a down-regulation of cell cycle genes^12,17,19^, however, these experiments were done in a non-acute setting. The fact that it takes more than 24 hours after depletion to see the first effects on cell proliferation seems to suggest that ZFP143 is not directly involved in the regulation of the cell cycle. Rather, the proliferation defect may be more consistent with being a consequence of the mitochondrial phenotype, which may result in reduced energy production^100,105–108^. In this case, ZFP143 contributes positively to proliferation by regulating genes involved in energy homeostasis.

Our data are consistent with existing literature suggesting that ZNF143 is crucial for development. Knockdown of ZFP143 in mESCs results in loss of pluripotency, suggesting that ZFP143 is required to maintain the pluripotent state^23^. Although our results do not directly implicate ZFP143 in regulating pluripotency genes, we observed an up-regulation of differentiation genes after 24 hours of ZFP143 depletion. However, acute depletion of *bona fide* pluripotency transcription factors such as OCT4 or SOX2^109,110^ leads to rapid changes in morphology and differentiation, which we did not observe for ZFP143 depletion. Paradoxically, loss of ZFP143 in the early stages of gastruloid development results in a failure to differentiate. The majority of differentiated cell types begin to form between 72 and 96 hours of gastruloid development, and cells in gastruloids lacking ZFP143 from 48 hours onwards fail to progress beyond the progenitor state. Since ZFP143-deficient cells can exit the pluripotent state upon retinoic acid treatment, it seems more likely that mitochondrial homeostasis maintained by ZFP143 transcriptional activation is critical for gastruloid development. In line with our finding that depletion of ZFP143 for longer than 24 hours causes a reduction in cell proliferation and mitochondrial dysfunction, gastruloids deficient for ZFP143 from 48 hours onwards should exhibit these phenotypes within 72-96 hours window^111–113^. The absence of this effect at later treatment times may be explained by the fact that the gastruloids have already undergone the necessary molecular and metabolic changes^114,115^. Nevertheless, the later onset of mitochondrial dysfunction and proliferation defects is manifested by the morphological changes. Our results indicate that in a multicellular embryo model, as opposed to a cell line model, differentiation, defined as a change in cellular state, requires functional mitochondria.

Our data place ZNF143 at the centre of a gene regulatory network that controls mitochondrial function. By maintaining gene expression of a subset of nuclear-encoded mitochondrial genes, ZNF143 plays a critical role in maintaining cellular homeostasis, which may explain its previously identified roles in a variety of biological processes in normal and malignant conditions. We expect that our study will bring the focus back to the function of ZNF143 as an essential transcriptional activator, as it was originally described.

## Acknowledgements

We thank NKI Flow Cytometry Facility for help with single-cell sorting of genome-edited cells and flow cytometry analyses, NKI Genomics Core Facility for help with sequencing, NKI Proteomics Facility for mass-spectrometry data analysis, NKI BioImaging Facility for support of microscopy analysis, and NKI Research High Performance Computing for computational resources. We thank Anders Hansen and Domenic Narducci for the exchange of unpublished results. We thank Koen Flach for help with ZFP143-FKBP mESC line generation. We thank Moreno Martinović for help with 4C-seq data analysis. We thank Teun van den Brand for help with ATAC-seq data analysis.

We thank William Faller group at the NKI for fruitful discussions. We thank members of our lab for critically reading the manuscript. Work in the de Wit lab is supported by the Dutch Research Council (016.161.316, Vidi; VI.C.222.049, Vici) and the European Research Council (865459, “FuncDis3D”). The NKI Proteomics Facility is supported by the Dutch Research Council X-omics Initiative.

## Author contributions

M.D.M, M.M., and E.d.W. conceived and designed the study. M.M. engineered and characterised ZFP143-FKBP cell line. M.M. and H.T. prepared sequencing libraries. M.M., N.A.S., and H.T. performed functional and validation experiments. M.M., N.A.S., and L.B. performed embryo model experiments. M.D.M. analysed sequencing, mass spectrometry, flow cytometry, microscopy, and publicly available data. E.d.W. supervised the study. M.D.M., M.M. and E.d.W. wrote the manuscript with input from all authors. All authors read and approved the manuscript.

## Figure and table legends

**Figure S1.**
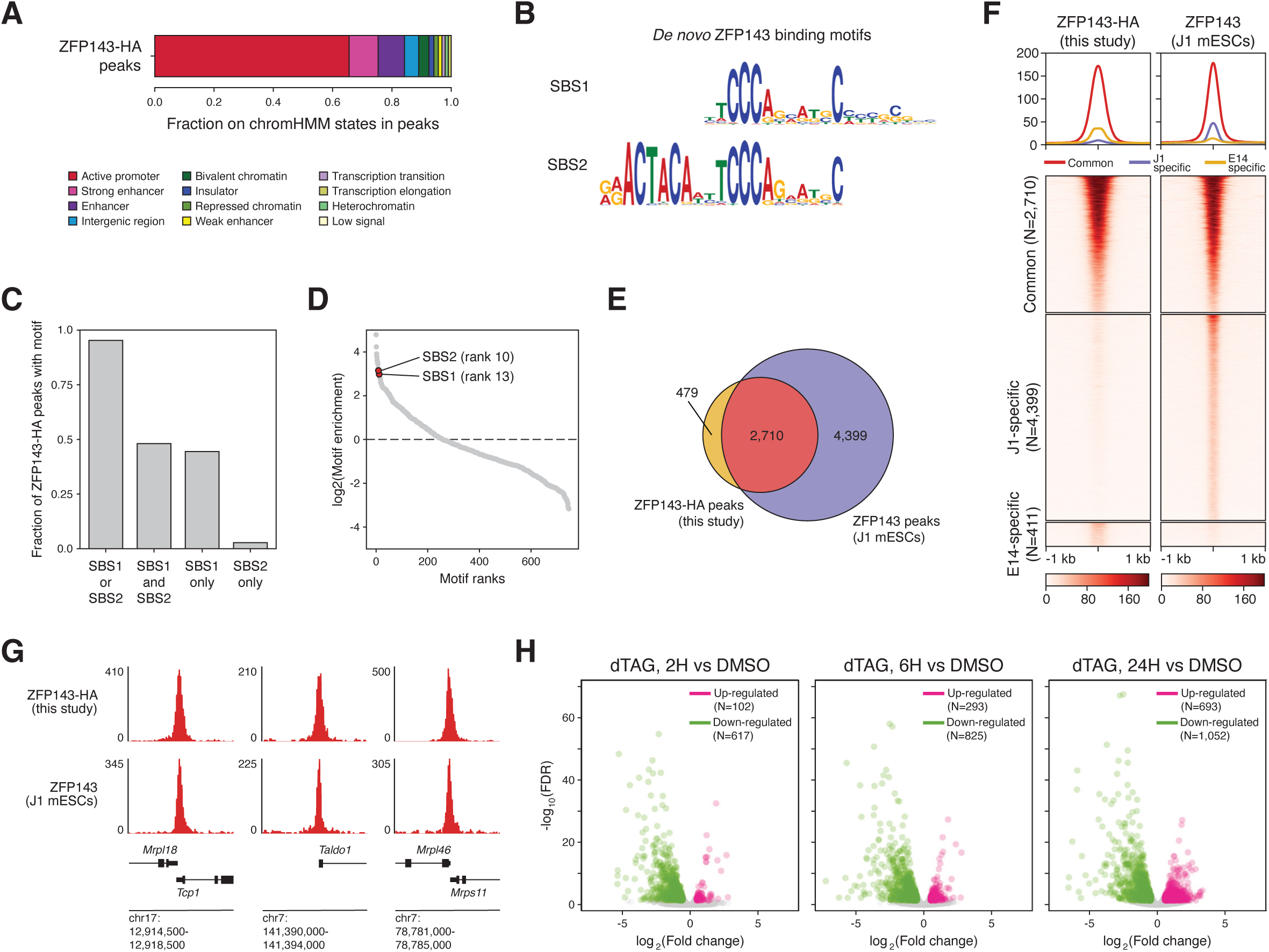
ChIP-seq and TT-seq in the ZFP143-FKBP cell line recapitulate known patterns of ZFP143 binding and gene regulation. **(A)** Distribution of ZFP143-HA ChIP-seq peaks across chromatin states annotated in mESCs^50^. Colours represent different chromatin states as in the original annotation. **(B)** *De novo* annotated SBS1 and SBS2 motifs in ZFP143-HA ChIP-seq peaks. **(C)** Fraction of ZFP143-HA peaks containing annotated SBS1 and SBS2 motifs. **(D)** Global motif enrichment analysis in ZFP143-HA peaks using the *de novo* annotated SBS motifs and motifs from the JASPAR database^59^. **(E)** Venn diagram showing the overlap between ZFP143-HA peaks and endogenous ZFP143 peaks detected in J1 mESCs with a custom antibody^12^ recognizing the endogenous ZFP143. **(F)** Tornado plots of ChIP-seq signals for ZFP143-HA in ZFP143-FKBP E14 mESCs and endogenous ZFP143 in J1 mESCs^12^ centred at common, J1-specific and E14-specific peaks. **(G)** Genomic tracks showing ChIP-seq signals of ZFP143-HA in ZFP143-FKBP E14 mESCs and endogenous ZFP143 in J1 mESCs^12^ at ZFP143 target loci, *Tcp1*, *Taldo1*, and *Mrps11*. **(H)** Volcano plots showing effect sizes and significance of the down-regulated (green) and upregulated (pink) genes measured by TT-seq in ZFP143-FKBP mESCs after 2, 6 and 24 hours of dTAG-V1 treatment, compared to DMSO treatment. The number of differentially expressed genes is indicated in the top right corners.

**Figure S2.**
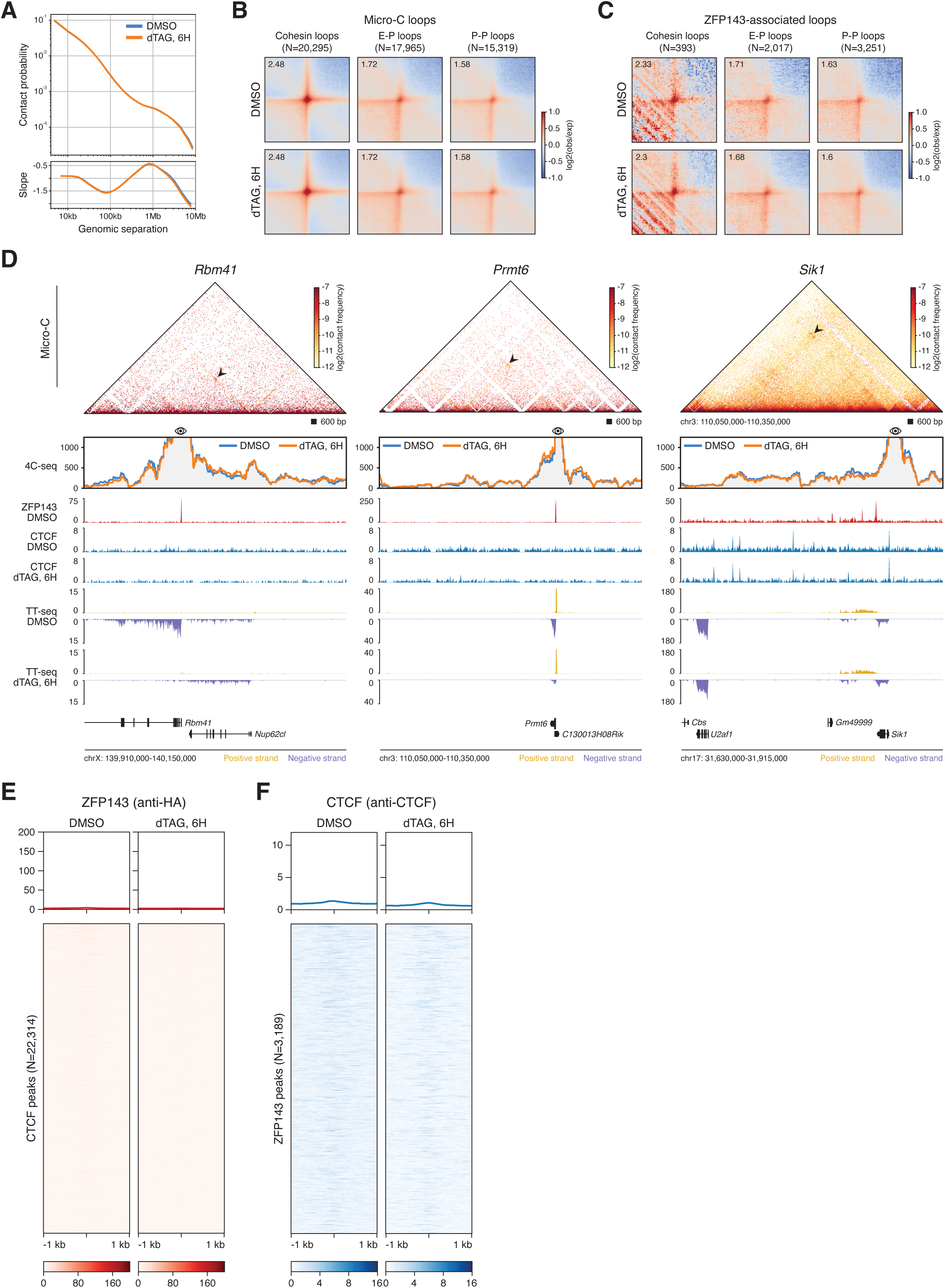
Chromatin interactions are largely unperturbed following ZFP143 depletion. **(A)** Relative contact probability plot (top panel) and its derivative (bottom panel) calculated from the Hi-C matrices of DMSO (blue) and 6 hours dTAG-V1 (orange) treated ZFP143-FKBP cells. **(B)** Average cohesin (left), enhancer-promoter (E-P, middle) and promoter-promoter (P-P, right) loops^54^ in DMSO and 6 hours dTAG-V1 treated ZFP143-FKBP cells. Value in the upper-right corner indicates the interaction strength of the loop over the background. **(C)** Same as in (B) but for the average ZFP143-associated loops (containing ZFP143 peak in at least one loop anchor). **(D)** High-resolution 4C-seq data generated for the ZFP143-bound genes *Rbm41* (left panel) and *Prmt6* (middle panel), and non-ZFP143-bound control gene *Sik1* (right panel), using gene promoters as viewpoints. The matrix in the top panel represents interaction frequencies in a previously published high-resolution Micro-C dataset^55^. The arrows point to detected Micro-C chromatin loops. The bottom panel shows 4C contact profiles in DMSO (blue) and in 6 hours dTAG-V1 (orange) treated ZFP143-FKBP cells. Genomic tracks show ZFP143-HA ChIP-seq (red), calibrated CTCF ChIP-seq (blue), TT-seq nascent transcription (yellow for sense and purple for antisense transcription) in control and 6 hours dTAG-V1 treated ZFP143-FKBP cells. **(E)** Tornado plots of ZFP143-HA ChIP-seq signal centred at CTCF peaks in DMSO and 6 hours dTAG-V1 treated ZFP143-FKBP cells. **(F)** Same as in (E) but for the calibrated CTCF ChIP-seq signal centred at ZFP143-HA peaks.

**Figure S3.**
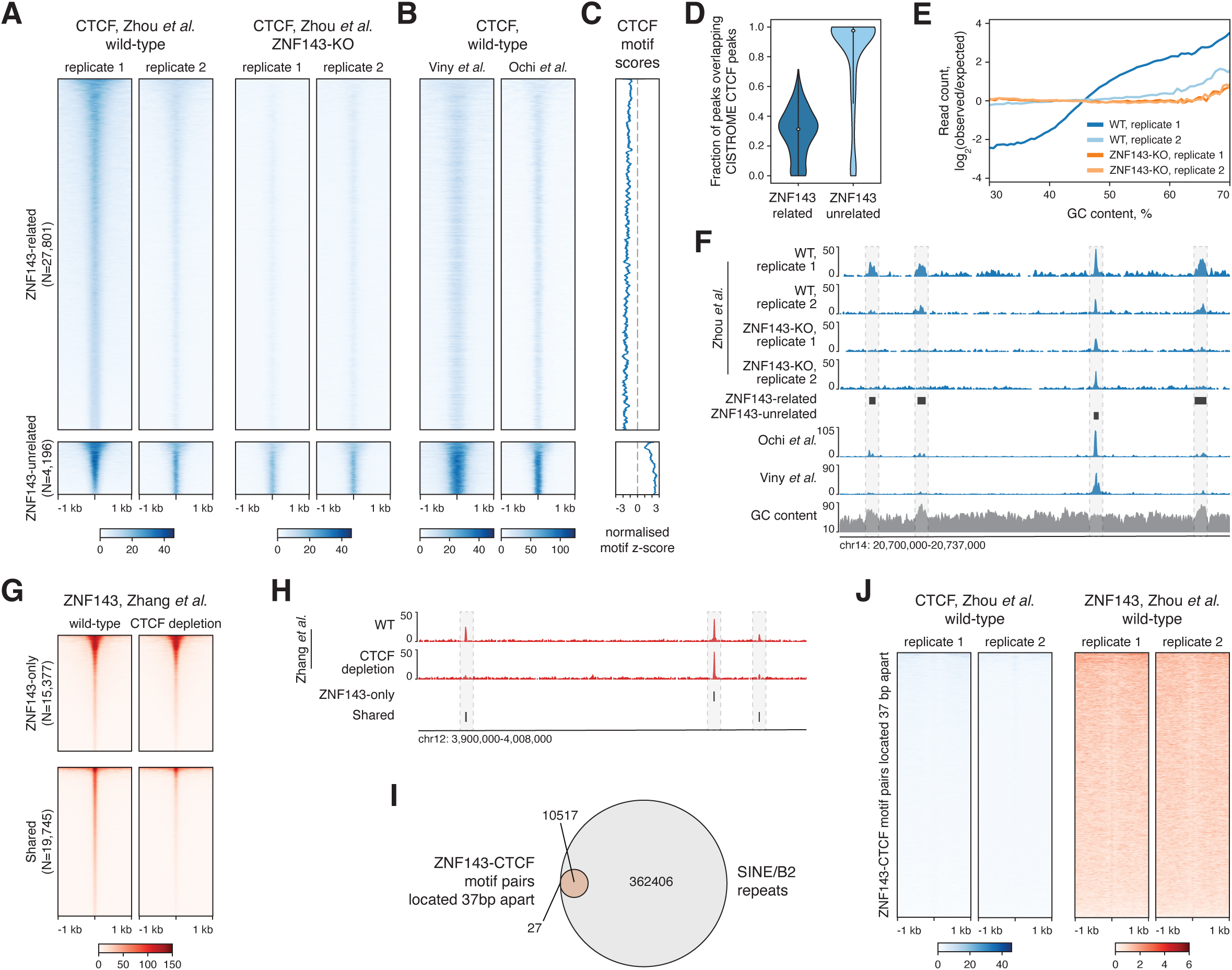
Re-analysis of previously published data suggests that CTCF-ZNF143 binding interdependency is likely an artefact. **(A)** Tornado plots of CTCF ChIP-seq signal from two biological replicates in wild-type (WT) and ZNF143-knockout (KO) haematopoietic stem and progenitor cells (HSPC) centred at ZNF143-related (top) and ZNF143-unrelated (bottom) CTCF peaks^38^. **(B)** Same as in (A) but for the CTCF ChIP-seq signal in HSPC from two orthogonal studies^117,118^. **(C)** Rolling mean of the normalised CTCF motifs scores, annotated for the ZNF143-related (top) and ZNF143-unrelated (bottom) CTCF peaks. **(D)** Violin plots showing the fraction of ZNF143-related (left) and ZNF143-unrelated (right) CTCF peaks overlapping CTCF peaks from the CISTROME database^57^. **(E)** GC bias scores calculated for CTCF ChIP-seq data generated from WT and ZNF143-KO HSPC samples^38^. Note the divergence of the first WT CTCF replicate from the rest of the samples. **(F)** Genomic tracks showing CTCF ChIP-seq signal from two biological replicates in WT and ZNF143-KO HSPC^38^, CTCF ChIP-seq signal from two other HSPC samples^117,118^, and GC content. Horizontal bars indicate ZNF143-related and ZNF143-unrelated CTCF peaks^38^. Note the overlap of ZNF143-related peaks with GC-rich regions. **(G)** Tornado plots of ZNF143 ChIP-nexus signal from control and CTCF-depleted HEC1B cells centred at ZNF143-only (top) and shared ZNF143 and CTCF (bottom) peaks^39^. **(H)** Genomic tracks showing ZNF143 ChIP-nexus signal from control and CTCF-depleted HEC1B cells^39^. Horizontal bars indicate ZNF143-only and shared ZNF143 and CTCF peaks. Note the specific loss of signal at shared peaks upon CTCF depletion. **(I)** Venn diagram showing the overlap between ZNF143-CTCF motif pairs located 37 bp apart from each other^38^ and SINE/B2 repeat elements in the mouse genome from RepeatMasker. **(J)** Tornado plots of CTCF and ZNF143 ChIP-seq signal centred at ZNF143-CTCF motif pairs located 37 bp apart from each other^38^.

**Figure S4.**
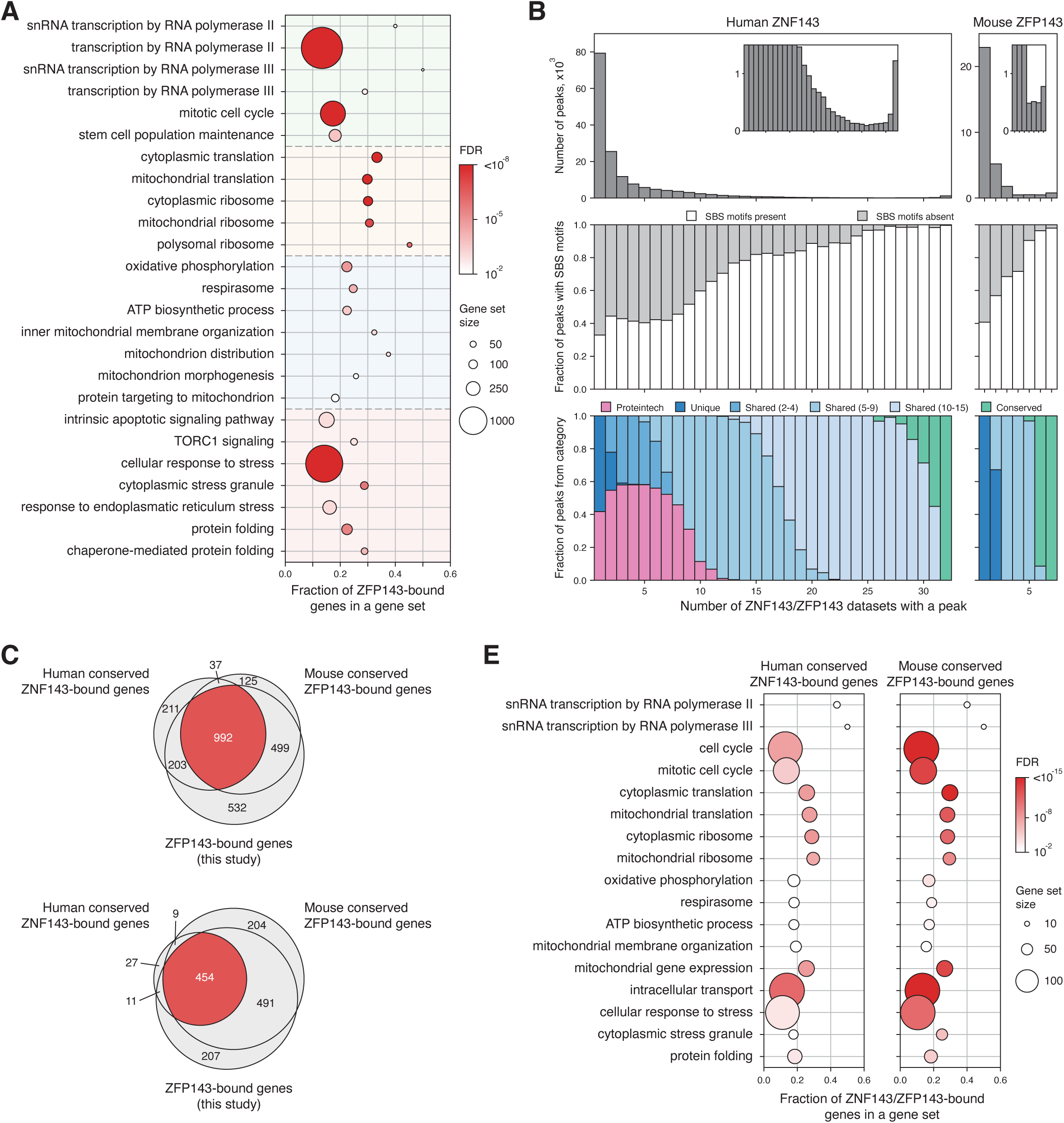
Conservation of ZFP143 targets across cell types and organisms. **(A)** Gene ontology (GO) terms, overrepresented in ZFP143-bound genes, identified based on ZFP143-HA ChIP-seq data in ZFP143-FKBP cells. Coloured areas represent gene sets involved in various cellular functions. **(B)** Bar plots showing the number of ZNF143/ZFP143 peaks overlapping between datasets (top panels), the fraction of ZNF143/ZFP143 peaks with SBS motifs present (middle panels), and the number of cell types sharing ZNF143/ZFP143 peaks (bottom panels) in the re-analysed publicly available human and mouse ChIP-seq datasets. **(C)** Venn diagram showing the overlap between conserved ZNF143-bound genes in human, conserved ZFP143-bound genes in mouse, and ZFP143-bound genes identified in ZFP143-FKBP cells. **(D)** Venn diagram showing the overlap between GO terms significantly overrepresented in conserved ZNF143-bound genes in human, conserved ZFP143-bound genes in mouse, and ZFP143-bound genes identified in ZFP143-FKBP cells. **(E)** GO terms, overrepresented in conserved ZNF143-bound genes in human (left panel) and conserved ZFP143-bound genes in mouse (right panel).

**Figure S5.**
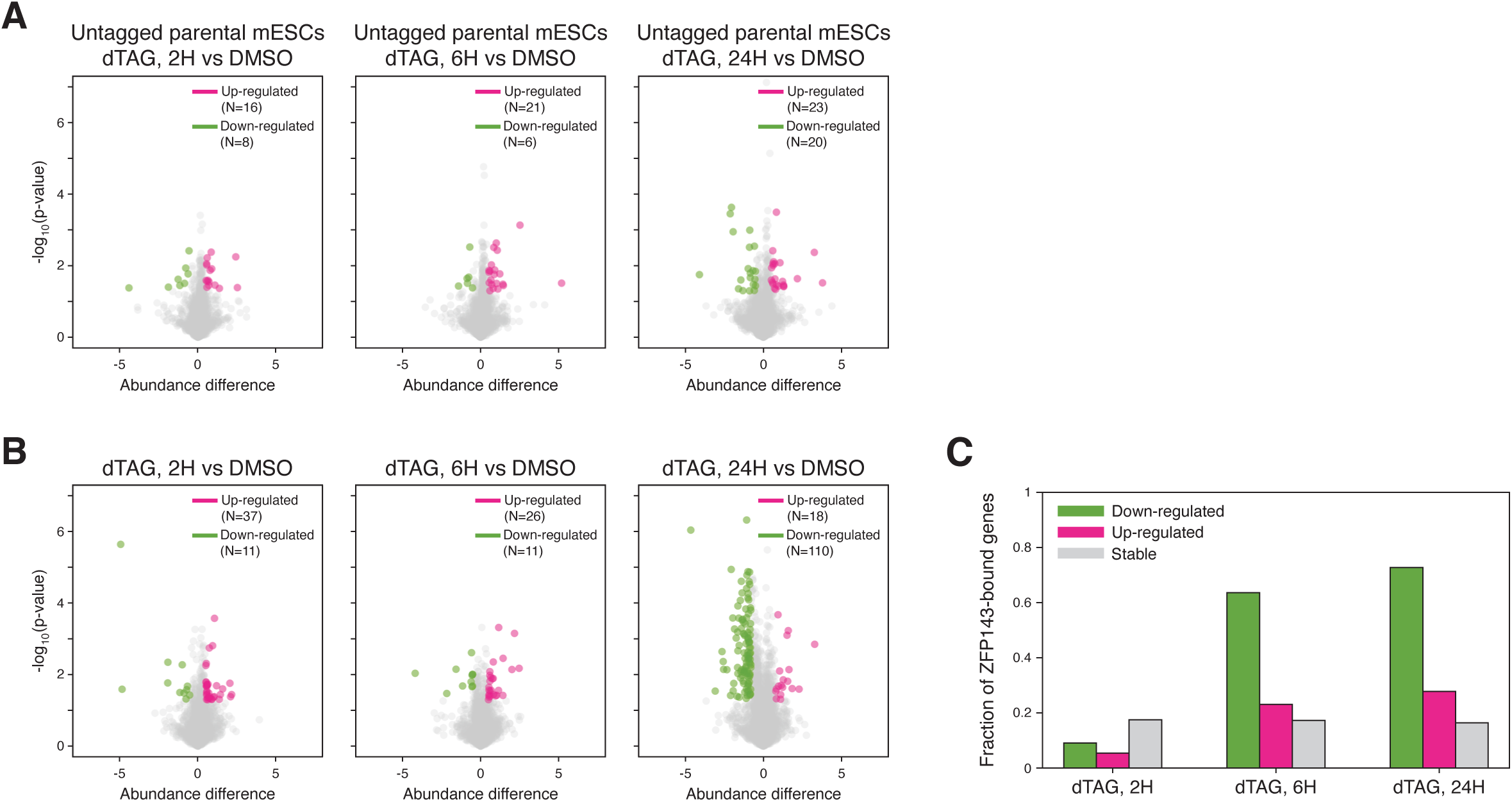
Proteome changes after ZFP143 depletion are delayed but consistent with changes in the nascent transcriptome. **(A)** Volcano plot showing effect sizes and significance of the down-regulated (green) and up-regulated (pink) proteins measured by quantitative mass spectrometry in untagged E14 mESCs after 2, 6 and 24 hours of dTAG-V1 treatment, compared to DMSO treatment. The number of differentially expressed proteins is indicated in the top right corners. **(B)** Same as (A) but for the ZFP143-FKBP cells. **(C)** Fraction of ZFP143-bound genes among down-regulated (green), up-regulated (pink), and stable (grey) proteins in the mass spectrometry at the indicated times after ZFP143 depletion.

**Figure S6.**
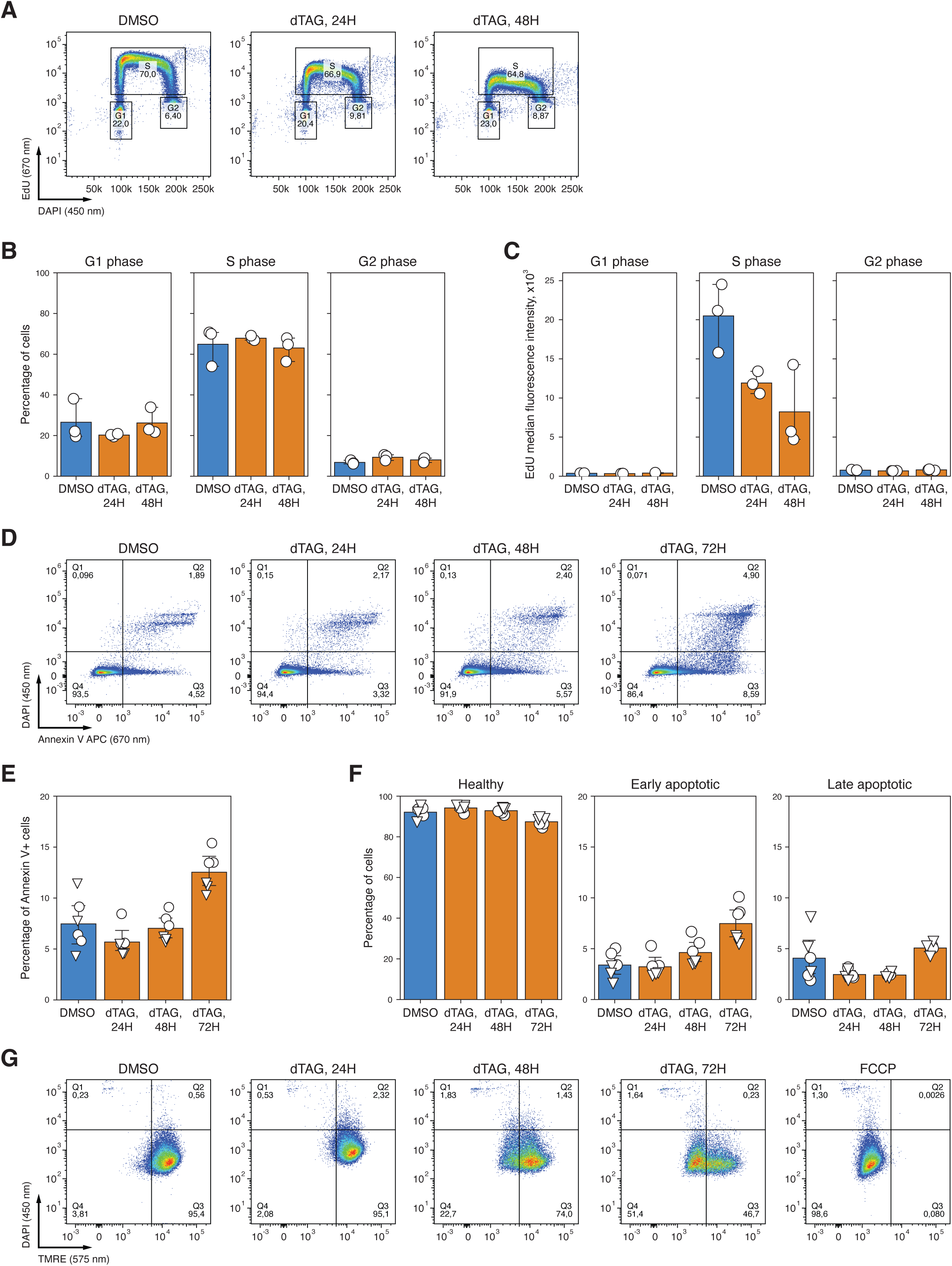
Flow cytometry analysis of cell cycle, apoptosis, and mitochondrial membrane potential following ZFP143 loss. **(A)** Representative flow cytometry data plots for EdU incorporation assay to measure cell cycle phases in DMSO, 24 and 48 hours dTAG-V1 treated ZFP143-FKBP cells. **(B)** Quantification of cell cycle phases in DMSO (blue) and 24 and 48 hours dTAG-V1 (orange) treated ZFP143-FKBP cells. Dots represent values for replicates. Error bars indicate 95% confidence interval. **(C)** Quantification of EdU median fluorescence intensity for different cell cycle phases in DMSO (blue) and 24 and 48 hours dTAG-V1 (orange) treated ZFP143-FKBP cells. Dots represent values for replicates. Error bars indicate 95% confidence interval. **(D)** Representative flow cytometry data plots for Annexin V staining to measure apoptosis rates in DMSO, and 24, 48 and 72 hours dTAG-V1 treated ZFP143-FKBP cells. **(E)** Quantification of total Annexin V-positive cells in DMSO (blue), and 24, 48 and 72 hours dTAG-V1 (orange) treated ZFP143-FKBP cells. Dots represent values for replicates. Error bars indicate 95% confidence interval. **(F)** Quantification of healthy, early apoptotic and late apoptotic cells in DMSO (blue), and 24, 48 and 72 hours dTAG-V1 (orange) treated ZFP143-FKBP cells. Dots represent values for replicates. Error bars indicate 95% confidence interval. **(G)** Representative flow cytometry data plots for TMRE staining to measure mitochondrial membrane potential in DMSO, 24, 48 and 72 hours dTAG-V1, and FCCP treated ZFP143-FKBP cells.

**Figure S7.**
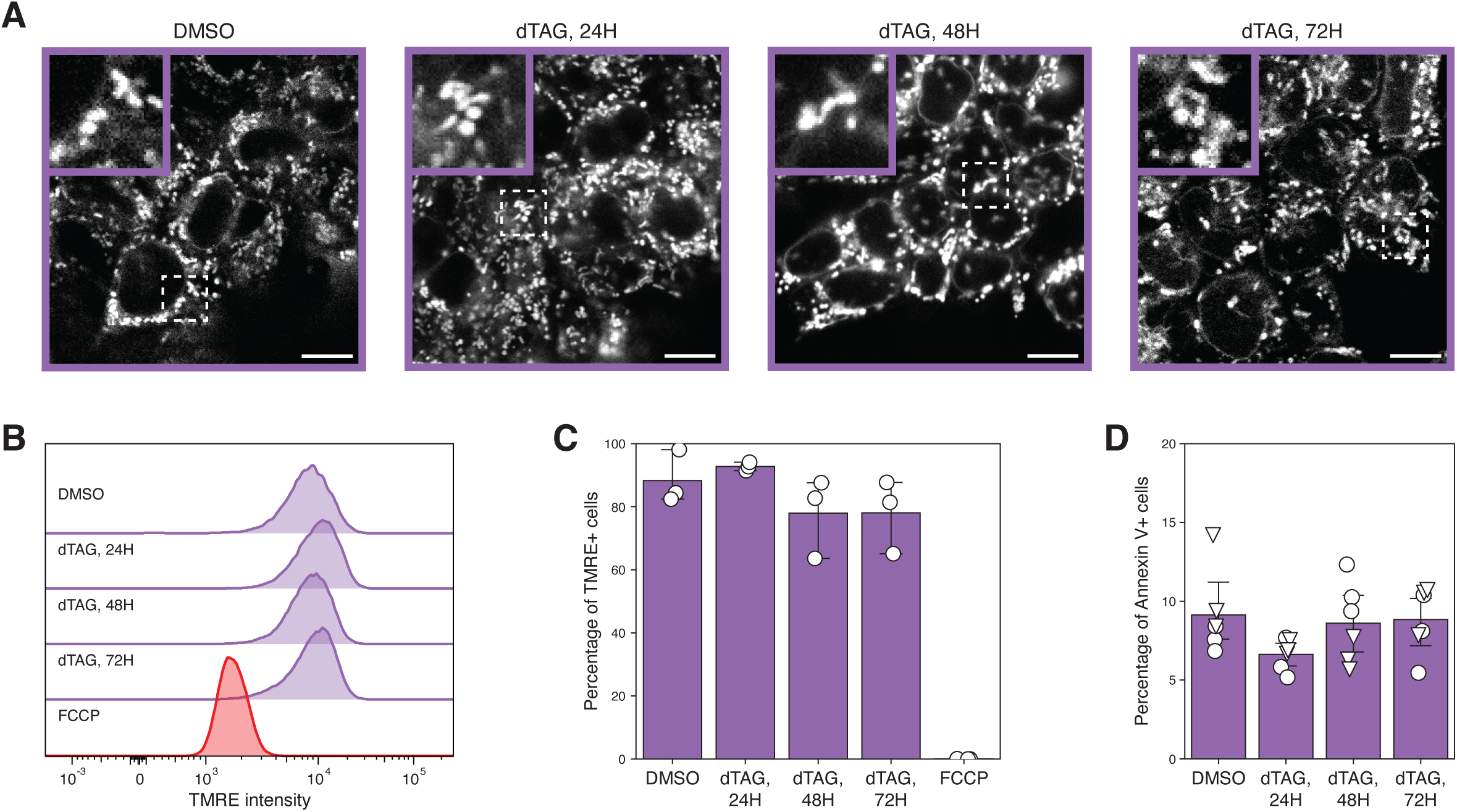
Treatment of untagged E14 mESCs with dTAG-V1 does not affect mitochondria. **(A)** Representative confocal microscopy images of the MitoTracker Red FM fluorescence in DMSO and 24, 48, and 72 hours dTAG-V1 treated untagged E14 mESCs. Magnified images of mitochondrial morphology and mitochondrial network for the dash-boxed regions in the representative images are shown in the upper-left corners. Scale bar: 10 µm. **(B)** Representative flow cytometry histogram showing TMRE fluorescence distribution in DMSO (purple), 24, 48, and 72 hours dTAG-V1 (purple), and FCCP (red) treated untagged E14 mESCs. **(C)** Quantification of TMRE-positive cells in DMSO (purple), 24, 48 and 72 hours dTAG-V1 (purple), and FCCP (red) treated untagged E14 mESCs. Dots represent values for replicates. Error bars indicate 95% confidence interval. **(D)** Quantification of total Annexin V-positive cells in DMSO and 24, 48 and 72 hours dTAG-V1 treated ZFP143-FKBP untagged E14 mESCs. Dots represent values for replicates. Error bars indicate 95% confidence interval.

**Figure S8.**
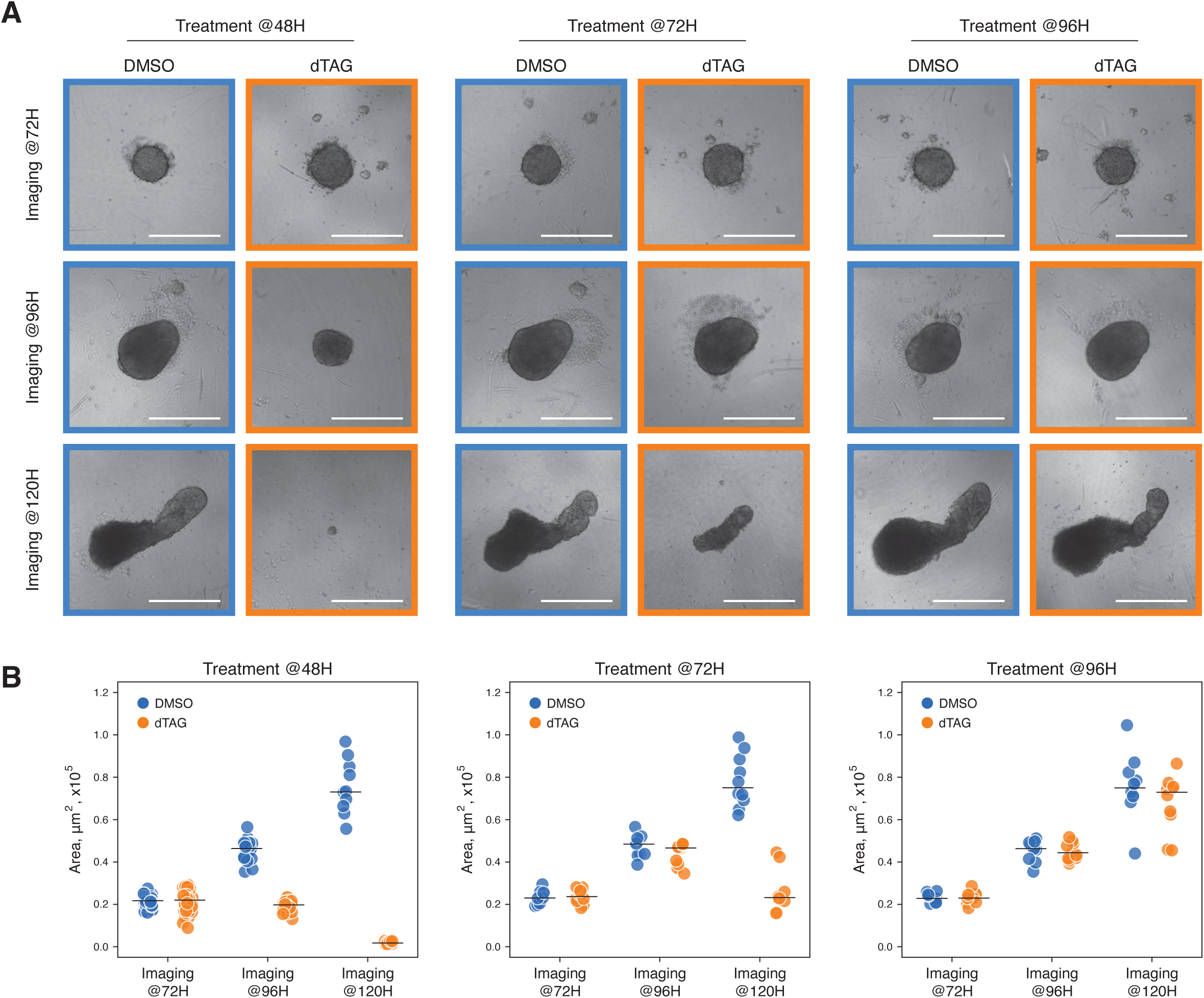
Time-course imaging and analysis of gastruloids morphology. **(A)** Bright-field microscopy images showing gastruloids treated with DMSO (blue frame) and dTAG-V1 (orange frames) at the times indicated on top and imaged at the developmental stages indicated on the left. Scale bar: 300 µm. **(B)** Quantification of gastruloid sizes for gastruloids treated with DMSO (blue) and dTAG-V1 (orange) from (A). Black lines indicate the mean values. The number of quantified gastruloids is indicated below.

**Figure S9.**
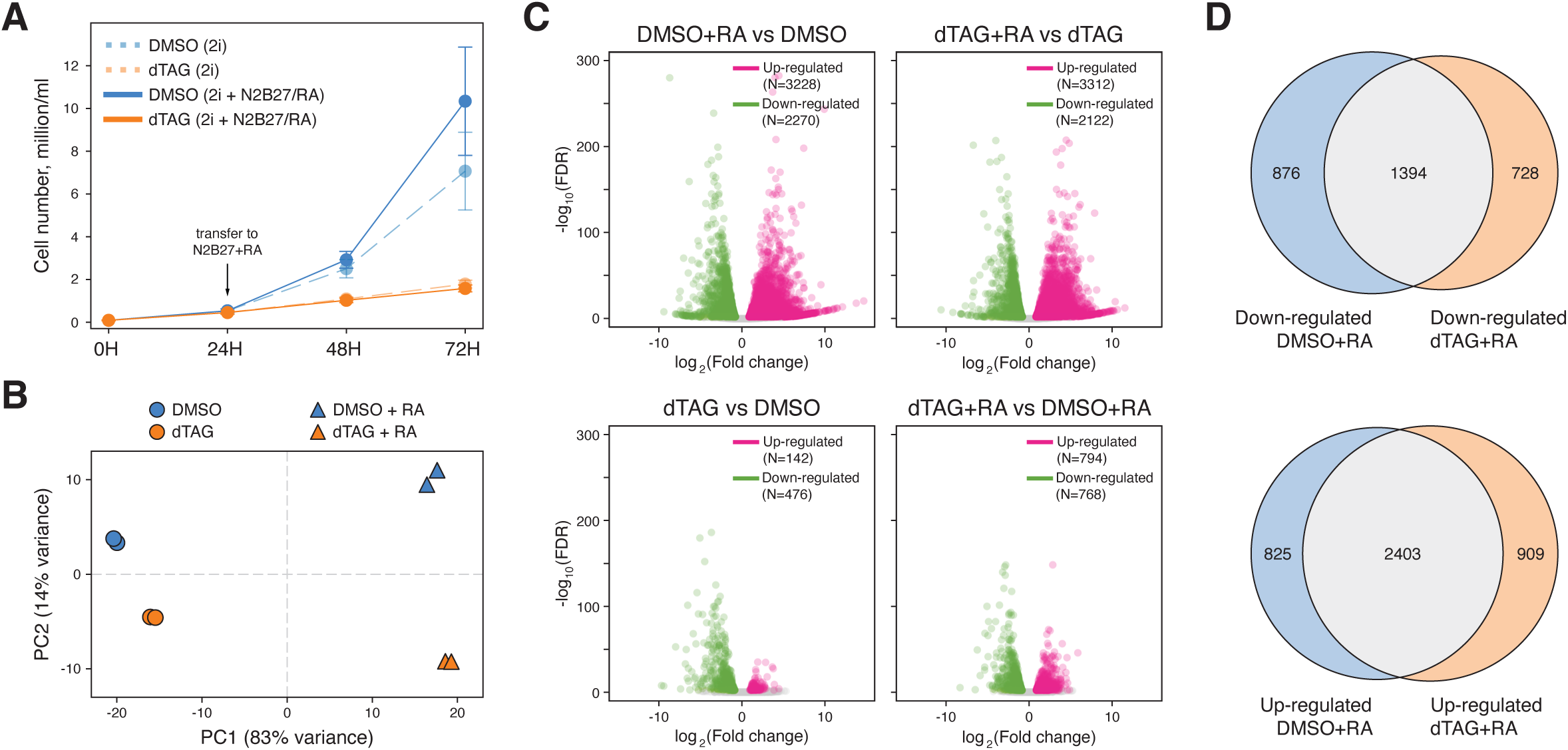
Transcriptomic changes upon differentiation with retinoic acid. **(A)** Growth curve showing the total cell number of DMSO (blue) and dTAG-V1 (orange) treated ZFP143-FKBP cells grown in 2i media (dashed line) and after transfer to N2B27 differentiation media with retinoic acid 24 hours post-depletion start (solid line). Dots indicate mean values. Error bars indicate standard deviation. **(B)** Principal component analysis of the RNA-seq data of DMSO (blue) and dTAG-V1 (orange) treated ZFP143-FKBP cells grown in 2i media (circles) and after transfer to N2B27 differentiation media with retinoic acid 24 hours post-depletion start (triangles). **(C)** Volcano plot showing effect sizes and significance of the down-regulated (green) and upregulated (pink) genes measured by RNA-seq before and after retinoic acid treatment in DMSO and dTAG-V1 treated ZFP143-FKBP cells. Top row shows volcano plots for the effect of retinoic acid in DMSO (left panel) and dTAG-V1 (right panel) treated cells. Bottom row shows volcano plots for the effect of dTAG-V1 treatment before (left panel) and after (right panel) transfer to differentiation media. The number of differentially expressed genes is indicated in the top right corners. **(D)** Venn diagram showing the overlap between down-regulated (top panel) and up-regulated (bottom panel) genes in DMSO and dTAG-V1 treated mESCs before and after differentiation with retinoic acid.

**Table S1. List of the oligonucleotides and primers used in this study.**

**Table S2. List of the publicly available datasets re-analysed in this study.**

**Table S3. Overlap between re-analysed ZNF143 peaks and CISTROME CTCF peaks.**

**Table S4. List and gene ontology analysis of the ZFP143/ZNF143-bound genes.**

**Table S5. Differential gene expression of the TT-seq data.**

**Table S6. Annotation of the differentially expressed genes with MitoCarta database.**

**Table S7. Gene set enrichment analysis of the TT-seq data.**

**Table S8. Differential protein abundance of the mass spectrometry data.**

**Table S9. Gene set enrichment analysis of the mass spectrometry data.**

**Table S10. Differential gene expression of the RNA-seq data.**

**Table S11. Gene set enrichment analysis of the RNA-seq data.**

## STAR Methods

### Resource availability

#### Lead contact

Further information and requests for resources and reagents should be directed to and will be fulfilled by the lead contact, Elzo de Wit.

#### Materials availability

Plasmids generated in this study have been deposited to Addgene and will be available upon publication in a journal. ZFP143-FKBP mESC cell line generated in this study is available upon reasonable request from the lead contact, Elzo de Wit.

#### Data and code availability

● ChIP-seq, Hi-C, 4C-seq, TT-seq, and RNA-seq data have been deposited at the GEO database (GSE260914) and are publicly available as of the date of publication. Proteomics data have been deposited at the PRIDE database (PXD049480) and are publicly available as of the date of publication. Previously published ChIP-seq datasets used in this study can be accessed from the GEO database. All accession numbers are listed in the key resources table.
● All original code has been deposited at Github (https://github.com/magnitov/zfp143) and is publicly available as of the date of publication.
● Any additional information required to reanalyze the data reported in this paper is available from the lead contact upon request.

### Experimental model and study participant details

#### Cell lines

Mouse Embryonic Stem Cells E14Tg2A (129/Ola) cell lines were cultured on 0.1% gelatin-coated plates in serum-free DMEM/F12 (Gibco) and Neurobasal (Gibco) medium (1:1) supplemented with N-2 (Gibco), B-27 (Gibco), BSA (0.05%, Gibco), 10 × 4 U of Leukaemia Inhibitory Factor (LIF) (Millipore), MEK inhibitor PD0325901 (1 μM, Selleckchem), GSK3-β inhibitor CHIR99021 (3 μM, Cayman Chemical) and 1-Thioglycerol (1.5 × 10^−4^ M, Sigma-Aldrich). The cell lines were passaged every 2 days in daily culture. For treatment using small molecule degraders, cells were counted and seeded as normal. The day after seeding DMSO or dTAG-V1 at a final concentration of 500nM (in DMSO) were added to cell medium. Molecules and media were refreshed every 24h if needed. Cells were tested for Mycoplasma routinely (Lonza).

#### Gene targeting

For the knock-in of the FKBP sequence at the genes of interest, previously described approach^119^ and plasmids to recognize the endogenous Zfp143 locus were used (ZFP143-sgRNA and ZFP143-FKBP-donor, will be available through Addgene upon publication in a journal). The DNA sequence for the knock-in was designed to include the FKBP-2xHA-P2A-GFP cassette in between two homology arms for surrounding the STOP codon of *Znf143* gene and ordered from Twist Bioscience. Plasmids were amplified in competent bacteria and prepared for nucleofection following Nucleofector kit protocol (Lonza). The cells were transfected with the plasmids containing the gRNA sequence and the donor plasmid designed to include the FKBP-2xHA-P2A-GFP in between two homology arms for the gene of interest. After transfection, cells were grown in 2i medium. After expansion, single cells with a positive GFP signal were sorted in 96-well plates and manually picked for genotyping using PCR and western blotting. Homozygous clones responding to dTAG-V1 were used for experiments. Primers and gRNA sequences are listed in Table S1.

### Method details

#### Cell count

For cell count, 4×10^4^ ZFP143-FKBP cells were seeded. The following day the cells were counted to get 0 hour point and then DMSO or dTAG-V1 at a final concentration of 500 nM were added directly into the medium. Medium supplemented with treatment molecules was refreshed daily. For counting after 24, 48, and 72 hours or treatment, cells were harvested by trypsinization with TVP1X, gently dissociated into single cells and counted by an automated cell counter (Bio-Rad). For cell count during retinoic acid differentiation, the same procedure was followed, replacing the media with N2B27 medium supplemented with RA after 24 hours of pre-treatment with DMSO or dTAG-V1. All cell count experiments were performed in technical triplicates and repeated for two biological replicates.

#### Western blotting

Whole cell lysate was collected by adding RIPA Lysis buffer (150 mM NaCl, 1% NP-40, 0.5% Sodium Deoxycholate, 0.1%SDS, 25mM Tris (pH 7.4)) to the cells for 1 hour. Proteins were quantified using Bradford assays. 40µg of proteins were loaded and separated on an in-house prepared 10% SDS-PAGE gel. After transfer on a PVDF pre-activated membrane using the Trans-Blot turbo transfer system (Bio-Rad), membrane proteins were blocked with 5% milk. Blots were incubated with primary antibodies (HA, ab9110, Abcam, 1:2000; ZNF143, 16618-1-AP, Proteintech 1:1000; ɑ-Tubulin, T5168, Sigma-Aldrich, 1:5000) for 1 hour at room temperature. Species-specific secondary antibodies were incubated for 1 hour at room temperature. After incubation, blots were washed at least three times in 0.1% Tween-20 in TBS. Proteins were detected using the Clarity Western ECL Substrate (Bio-Rad) with ChemiDoc MP Imaging system (Bio-Rad).

#### ChIP-seq

For chromatin preparation, mESCs cells were grown and treated with 500 nM dTAG-V1 or DMSO for ZFP143 depletion. Cells were harvested and collected to have 1×10^6^ cells/ml. For CTCF ChIP-seq, 10% of HCT116 cells were added to mESC cells and used as an internal reference. Chromatin was cross-linked using 1% FA final concentration at RT for 10 min. Glycine 2.0 M was added directly after to quench the cross-linking reaction. Fixed cells were lysed and chromatin was sheared on a Bioruptor Plus sonication instrument (Diagenode) to achieve chromatin fragments between 200 and 500 bp. For the IP, antibodies were coupled overnight with Protein G Dynabeads (Thermo Scientific). The following day, antibodies-coupled beads were washed and the sheared chromatin was added. Antibodies against CTCF (07-729, Merck Millipore, 5 μl per ChIP) and HA (ab9110, Abcam, 10 μl per ChIP) were used. After incubation of the chromatin and antibody-beads overnight at 4°C, the chromatin fragments captured on beads were de-crosslinked from proteins and DNA fragments were released. DNA was purified using the MiniElute PCR purification kit (Qiagen). The resulting DNA was used for preparation of the sequencing library, following the KAPA HTP Library Preparation kit (Roche). Libraries were sequenced on Illumina NextSeq 550 generating paired-end reads.

#### Hi-C

Hi-C data was generated as previously described^40^ with minor modifications^81^. For each template, 10 million cells were harvested and crosslinked using 2% formaldehyde. Crosslinked DNA was digested in the nucleus using MboI (NEB), and biotinylated nucleotides were incorporated at the restriction overhangs and joined by blunt-end ligation. The ligated DNA was enriched in a streptavidin pull-down. Hi-C libraries were prepared using a standard end-repair and A-tailing method and sequenced on Illumina MiSeq and NextSeq 550 generating paired-end reads.

#### 4C-seq

4C was performed for DMSO and 500 nM dTAG-V1 treated ZFP143-FKBP cells as previously described^56,120^ using a two-step PCR method for indexing^81^. 10 million cells were used for each time-point. MboI (NEB) was used as the first restriction enzyme and Csp6I (NEB) as the second restriction enzyme. Viewpoint-specific primers are listed in Table S1. 4C-seq was performed in two biological replicates. Libraries were sequenced on Illumina NextSeq 550 generating paired-end reads.

#### TT_chem_-seq

Libraries for TT_chem_-seq were prepared following a published protocol^52^. ZFP143-FKBP cells were seeded and the day after treated with 500 nM of DMSO or dTAG-V1 for 2, 6, and 24 hours. Cells were labelled with 2 mM of the uridine analog 4-thiouridine (Sigma-Aldrich) for 10 min. Total RNA was isolated and fragmented. The 4sU-biotin labelled RNA was enriched using µMacs Streptavidin Kit (Miltenyi). Libraries were prepared using KAPA RNA HyperPrep and KAPA Dual-Indexed Adapter kits (Roche) using dual indexing adapters. TT_chem_-seq was performed in two biological replicates. Libraries were sequenced on Illumina NextSeq 550 generating paired-end reads.

#### Quantitative mass spectrometry

For quantitative measure of protein abundance, ZFP143-FKBP and untagged parental mESCs were expanded and treated with either DMSO or 500 nM dTAG-V1 for 2, 6 and 24 hours. Cells were harvested and 30 million cells per condition were centrifuged at 500g for 5 minutes, washed with PBS and pellets were snap frozen. All samples were prepared in triplicate.

For protein digestion, frozen pellets were lysed in boiling Guanidine (GuHCl) lysis buffer as previously described^121^. Protein concentration was determined with a Pierce Coomassie (Bradford) Protein Assay kit (Thermo Scientific), according to the manufacturer’s instructions. After dilution to 2M GuHCl, aliquots corresponding to 200µg of protein were digested twice (4h and overnight) with trypsin (Sigma-Aldrich) at 37°C, enzyme/substrate ratio 1:75. Digestion was quenched by the addition of FA (final concentration 5%), after which the peptides were desalted on a Sep-Pak C18 cartridge (Waters) and dried in a vacuum centrifuge. Prior to mass spectrometry analysis, the peptides were reconstituted in 2% formic acid. Peptide mixtures were analysed by nano LC-MS/MS on an Orbitrap Exploris 480 Mass Spectrometer equipped with an EASY-nLC 1200 system (Thermo Scientific). Samples were directly loaded onto the ReproSil-Pur 120 C18-AQ analytical column (2.4 μm, 75 μm × 500 mm, packed in-house). Solvent A was 0.1% formic acid/water and solvent B was 0.1% formic acid/80% acetonitrile. Samples were eluted from the analytical column at a constant flow of 250 nl/min in a 90-min gradient, containing a 78-min linear increase from 6% to 30% solvent B, followed by a 12-min wash at 90% solvent B.

#### Flow cytometry

##### Cell cycle assay

Cell cycle analysis was performed following the protocol of the Click-iT EdU Alexa Fluor 647 Flow Cytometry Assay kit (Invitrogen). Briefly, cells were counted, seeded and treated with either DMSO or dTAG-V1 the following day. For labelling, 10 uM EdU was added to cells, for 1.5 h and incubated at 37°C. After this time cells were harvested, fixed and permeabilized using a saponin-based permeabilization and wash reagents provided from the manufacturer. To detect EdU, cells were incubated with Click-iT reaction cocktail for 30 min. For DNA content measure, RNA was removed adding 10 μg/ml Ribonuclease A and DAPI (1:1000) was added to stain for DNA content. EdU and DAPI fluorescent signals were quantified on a BD LSRFortessa analyzer (BD Biosciences) with the R(C) 670/30 and V(F) 450/50 laser settings. Cell count assay was performed in technical triplicates and repeated for one biological replicate. Subsequent analysis was done with FlowJo v10.9.0 software. Single cells were gated, and G1, S and G2 cells were separated and quantified using the indicated gates.

##### Apoptosis assay

Apoptosis detection was performed by using Annexin V APC (BioLegend). After harvesting, the cells were resuspended in 150 μl binding buffer (0.1 M HEPES pH 7.4, 1.4 M NaCl, 25 mM CaCl2) and spinned down. The supernatant was discarded and 300 μl of Annexin V Binding Buffer, 3 μl of Annexin V APC (stock concentration 8μg/ml) and 3 μl DAPI (stock concentration 70μg/ml) were added to the cells. The cells were incubated for 20 minutes at room temperature and Annexin V and DAPI fluorescent signals were then quantified on a BD FACSAria Fusion analyzer (BD Biosciences) with the R(C) 670/30 and V(F) 450/50 laser settings. Apoptosis assay was performed in technical triplicates and repeated for two biological replicates. Subsequent analysis was done with FlowJo v10.9.0 software. All single cells were gated, and healthy, early apoptotic and late apoptotic cells were then separated and quantified using the indicated gates.

##### Mitochondrial membrane potential assay

Mitochondrial membrane potential was measured by using TMRE (Thermo Scientific). Cells were trypsinized, washed with PBS and made single cells in suspension. As a control, 20 µM carbonyl cyanide 4-(trifluoromethoxy)phenylhydrazone (FCCP) was added to one of the samples and incubated for 30 minutes. Next, cells were incubated with 100 nM TMRE (Thermo Scientific) for 30 minutes. DAPI at working concentration 1µg/ml was added to the cells 15 mins prior to sorting. TMRE and DAPI signals were detected on a BD LSRFortessa analyzer (BD Biosciences) with the YG(E) 585/15 and V(F) 450/50 laser settings. Mitochondrial membrane potential assay was performed in technical triplicates and repeated for one biological replicate. Subsequent analysis was done with FlowJo v10.9.0 software. Single cells were gated, and TMRE-positive cells were then separated and quantified using the indicated gates.

#### Live-cell imaging of mitochondria

ZFP143-FKBP and untagged parental E14 mESC were pre-cultured in phenol free media in a 8-well chamber (ibidi) pre-coated with biolaminin (BioLamina). The 5×10^3^ cells were seeded per well and DMSO or 500 nM dTAG-V1 treatments were added to the appropriate wells. Prior to imaging, the media was removed and cells were incubated with 100nM of Mitotracker Red FM (Thermo Scientific) in culture medium with DMSO or dTAG-V1 for 30 min. After that, the staining solution was replaced with culture medium and the corresponding treatment. Live imaging was performed on a SP8 Leica confocal microscope. Mitochondria were imaged under a 63×/1.40 oil HC PL APO CS2 objective. Mitotracker Red FM was excited with a white light laser set to 595 nm and emission light was collected with a HyD SMD detector set from 640 to 730 nm. Live-cell imaging of mitochondria in ZFP143-FKBP cells was performed for two biological replicates.

#### Gastruloids

Gastruloids were cultured following a previously established protocol^122^. In brief, the formation of gastruloids involved the aggregation of mESCs in differentiation medium (N2B27). This medium comprises a combination of DMEM/F12 (Thermo Scientific) and Neurobasal medium (Thermo Scientific), supplemented with 0.5X N2 (Thermo Scientific), 0.5X B27 with retinoic acid (Thermo Scientific), 2mΜ Glutamine, β-mercaptoethanol, and 50U/ml penicillin-streptomycin. U-bottom 96-well plates (Thermo Scientific) were utilised for the culture, and the process was carried out in a humidified incubator set at 5% CO2 and 37°C. Cell dissociation was performed in N2B27 medium using a serological pipette paired with a 200µL pipette tip. Subsequently, 3.75×104 cells were transferred into N2B27 medium, resulting in a final volume of 5ml. A 40µL aliquot of this suspension, containing approximately 300 cells, was added to each well. At the 48-hour mark, 150µL of N2B27 medium with 3μM CHIR99021 was introduced using the Hamilton Star R&D liquid handling platform. Starting from the following day, the culture medium was replaced with fresh N2B27 medium on a daily basis.

##### Imaging

Image documentation was carried out using an inverted wide-field microscope (Zeiss Axio Observer Z1 Live) at 72, 96, and 120 hours time points. Images for DMSO and dTAG-V1 treatment at 120 hours were collected for three biological replicates. Images for DMSO and dTAG-V1 time-course treatment were collected for one biological replicate.

##### ATAC-seq

The Tn5 protein was purified by the NKI protein facility as previously described^74^. To achieve transposon annealing, 10 µL of 10X TE buffer was combined with 45 µL of 100µM Tn5MErev oligonucleotides and 45 µL of 100µM corresponding Tn5ME-A and Tn5ME-B oligonucleotides (listed in Table S1). Subsequently, the adapter solution underwent incubation at 95°C for 10 minutes and was gradually cooled to 4°C at a rate of 0.1°C per second. For transposome formation, the annealed adapters were diluted with an equal volume of H2O. This resulting adapter solution was then mixed with 0.2mg/mL Tn5 at a 1:20 ratio and incubated for 1 hour at 37°C. The transposomes were either directly utilised for tagmentation or stored at -20°C for a maximum of 2 weeks.

Next, gastruloids were collected and dissociated to a single cell suspension by gentle trypsinization. A total of 50,000 cells were gathered in cold PBS and subjected to lysis using a 2x lysis buffer (1 M Tris–HCl pH 7.5, 5 M NaCl, 1 M MgCl2, 10% IGEPAL). Subsequently, the cells were centrifuged, and the resulting pellet was treated with 2xTD buffer (20 mM Tris(hydroxymethyl)aminomethane; 10 mM MgCl2, 20% dimethylformamide, brought at pH 7.6 with acetic acid) and 2 uL of transposon mix. Following this, PCR amplification was conducted twice using KAPA HiFi HotStart PCR ReadyMix (Roche) and P5 and P7 indexed primers (refer to the oligonucleotide list). Fragments ranging from 200 to 700 bp were purified utilising AMPure XP beads (Beckman Coulter), and the quality of the DNA was assessed through Bioanalyzer High Sensitivity DNA analysis (Agilent).

#### Retinoic acid differentiation

For the differentiation using retinoic acid following ZFP143 depletion, ZFP143-FKBP cells were first seeded in normal 2i medium. One day after seeding, DMSO or dTAG-V1 at a final concentration of 500 nM were added to the cells for 24 hours. After that, cells were washed twice with PBS and differentiation medium (N2B27), containing DMEM/F12, supplemented with 0.5X N2 and 0.5X B27, 2mM Glutamine, β-mercaptoethanol and 50U/ml penicillin streptomycin supplemented with *all trans*-retinoic acid (R2625, Sigma-Aldrich) to 1µm final concentration (stock 1:1000 Sigma-Aldrich), was added to the cells. DMSO and dTAG were included to continue the depletion. For harvesting, the cells were dissociated using 2X TVP.

##### RNA-seq

For RNA-seq, cells were harvested, kept in an RLT buffer stored at -80°C until purification or were directly used for RNA purification. RNA was first isolated using standard RNA isolation procedure from Qiagen RNeasy Mini kit (Qiagen), including a DNaseI treatment. The resulting RNA was used directly for library preparation following the TruSeq Stranded mRNA kit (Illumina) with TruSeq RNA Single Indexes set A (Illumina). RNA-seq was performed in two biological replicates. Libraries were sequenced on Illumina NextSeq 550 generating paired-end reads.

### Quantification and statistical analysis

#### ChIP-seq

##### Data processing

ZFP143 ChIP-seq reads were mapped to the mm10 mouse reference genome assembly using bwa mem v0.7.17-r1188^123^. Calibrated CTCF ChIP-seq reads were mapped to the mm10 mouse and hg38 human reference genome assemblies using bwa mem v0.7.17-r1188^123^. Uniquely mapped reads with MAPQ > 10 mapped in proper read pairs (-f 2) were selected using SAMtools v1.12^124^. Duplicate reads were filtered out using the Picard v2.25.6 “MarkDuplicates” function.

Scaling factors for calibrated CTCF ChIP-seq normalisation were calculated as described previously^125^. First, to avoid potential bias, CTCF ChIP-seq peaks from the CISTROME database^57^ for E14Tg2A^60^ (ID: 85173, 85177) and HCT116^126^ (ID: 42148, 42149, 42150) were used, and only the peaks found in all biological replicates were extracted. Second, the number of reads covering peaks was calculated using the “multicov” function from BEDTools v2.27.1^127^, the immunoprecipitation efficiency was calculated as reads covering peaks divided by total number of reads for the mm10 and hg38 genomes, and the fraction of reads mapping to the spike-in hg38 genome was calculated. The absolute scaling factor for each sample was then calculated as the percentage of spike-in chromatin divided by the reads covering peaks for the spike-in genome and normalised by the immunoprecipitation ratio between spike-in and target genomes. Lastly, the relative scaling factors for all samples were calculated by dividing the obtained absolute scaling factors by the highest absolute scaling factor among all samples.

The bigwig coverage tracks for ZFP143 ChIP-seq were generated using the “bamCoverage” function from the deepTools v3.4.2^128^ with the “--effectiveGenomeSize” parameter set to 2652783500 and “--normalizeUsing” parameter set to RPGC. The bigwig coverage tracks for calibrated CTCF ChIP-seq were generated using the “bamCoverage” function from the deepTools v3.4.2^128^ and the calculated relative scaling factors specified under the “--scaleFactor” parameter.

##### Peak calling and analysis

Peak calling was performed using MACS2 v2.2.6^129^. For ZFP143 ChIP-seq, peaks were called in a narrowPeak mode using input as a control file, with mappable genome size set to mm, q-value cutoff of 0.01, and “--keep-dup” parameter set to all. For CTCF ChIP-seq, peaks were called in a narrowPeak mode using input as a control file, with mappable genome size set to mm, and q-value cutoff of 0.05. Only the peaks detected on the canonical chromosomes outside of the blacklist regions^130^ were retained. Since CTCF ChIP-seq demonstrated little difference between DMSO and dTAG treated conditions, the peaks detected in both cells were pooled together.

Pile-ups of the ChIP-seq signals were calculated using the “computeMatrix” function from deepTools v3.4.2^128^. Pile-ups were generated in reference-point mode with parameters “--referencePoint” center, “-a” 1000, “-b” 1000, “--missingDataAsZero”, and “--skipZeros”. The resulting matrices were visualised using plotHeatmap function from deepTools v3.4.2^128^. Genomic tracks plots were produced using pyGenomeTracks v3.8^131^.

*De novo* motifs discovery for ZFP143-HA peaks was performed using meme v5.4.1^132^. The sequences within the peaks were extracted using the “getfasta” function from BEDTools v2.27.1^127^. MEME was initialised with the parameters “-dna”, “-revcomp”, “-mod” zoops, “-nmotifs” 5 “-minw” 6, “-maxw” 50, “-markov_order” 0. Motif position weight matrices of the *de novo* discovered SBS1 and SBS2 motifs and core collection of JASPAR 2020 motifs in vertebrates^133^ were used to predict motif positions genome-wide with motifmatchr^134^. For this, “matchMotifs” function was used with background nucleotide frequencies set to “genome”. Motif enrichment in ZFP143-HA peaks was determined using the “permTest” function from regioneR^135^ by evaluating the number of overlaps between peaks and motifs. The background enrichment was assessed using 100 circular permutations of the peaks.

BEDTools v2.27.1 “intersect” function was used to calculate the overlaps between ZFP143-HA peaks and chromHMM states for mESC^50^. For comparison with publicly available ZFP143 data in mESC, ChIP-seq peaks from the CISTROME database^57^ for J1 mESC^12^ (ID: 36789) were used. The overlap between ZFP143-HA peaks and J1 mESCs peaks was calculated with BEDTools v2.27.1 “intersect” function.

#### Publicly available ChIP-seq data analysis

##### Re-mapping of publicly available ZNF143/ZFP143 datasets

For the systematic re-analysis of the ZNF143/ZFP143 binding, publicly available ZNF143/ZFP143 ChIP-seq datasets from human (N=32) and mouse (N=7) cells were collected (listed in Table S2). Where available, an appropriate input or IgG control was also re-analysed (listed in Table S2). ChIP-seq reads were mapped to the hg38/mm10 mouse reference genome assembly using bwa mem v0.7.17-r1188^123^. Due to the short read length of the IgG control samples, these data were mapped using bwa aln and bwa samse v0.7.17-r1188^123^. Uniquely mapped reads with MAPQ > 10 were selected using SAMtools v1.12^124^. For paired-end datasets, the reads were additionally required to be in proper read pairs (-f 2). Duplicate reads were filtered out using the Picard v2.25.6 “MarkDuplicates” function. Peak calling was performed using MACS2 v2.2.6^129^. All peaks were called in narrowPeak mode with input or IgG controls where applicable. MACS2 was run with mappable genome size set to hs or mm, q-value cutoff of 0.05, “--keep-dup” set to all, and a “--nomodel” parameter. For datasets where input or IgG control was present, the bigwig coverage tracks were generated using the “bamCompare” function from the deepTools v3.4.2^128^ with the “--operation” parameter set to ratio. For other datasets, the bigwig coverage tracks were generated using the “bamCoverage” function with the appropriate “--effectiveGenomeSize” parameter and “--normalizeUsing” set to RPGC.

##### Co-localization between ZNF143 and CTCF peaks

To analyse the co-localisation between ZNF143/ZFP143 peaks, uniformly processed publicly available CTCF ChIP-seq datasets from the CISTROME database^57^ were used. 512 CTCF peak sets for human and 460 peak sets for mouse were collected. For each ZNF143 dataset, the proportion of ZNF143 peaks overlapping with each CTCF peaks dataset was calculated using the "intersect" function from BEDTools v2.27.1^127^. Since CTCF binding is relatively conserved, the datasets were not matched according to the cell type. The resulting table showing the overlap between each re-analysed ZNF143/ZFP143 peak set and CTCF, as well as the accession numbers of the CTCF peak set in the CISTROME database, is provided in Table S3.

##### Re-analysis of ZNF143 ChIP-seq in K562 cell line

To explore the overlap between ZNF143 and CTCF peaks, ChIP-seq data generated in the K562 cell line with Proteintech and FLAG antibodies was used for comparison^58^. As both datasets had biological replicates, a list of reproducible peaks detected by Proteintech and FLAG antibodies within replicates was identified. The reproducible peak sets were then used to classify ZNF143 peaks into common, Proteintech-specific and FLAG-specific categories. Motif enrichment of *de novo* annotated SBS1/SBS2 and CTCF^59^ (JASPAR ID: MA0139.1) motifs in common, Proteintech-specific and FLAG-specific peaks was calculated with “permTest” function from regioneR^135^ as described above. To compare signals detected by different antibodies, the ZNF143 dataset produced with a custom antibody^12^ and the CTCF dataset from ENCODE^58^ with corresponding controls (see Table S2) were re-processed as described above. The pile-ups of the ChIP-seq signals were calculated using the “computeMatrix” function from deepTools v3.4.2^128^ as described above.

To reproduce the enrichment analysis at the loop anchors identified in the K562 cell line, the loop annotation from the original study was used^40^. In addition, peak datasets for 11 histone modifications, 144 transcription factors, and 1 DHS dataset from ENCODE used in the original publication were downloaded^40^ (listed in Table S3 of the original study). As loop annotation and peak sets were for the hg19 human reference genome assembly, the coordinates of the common, Proteintech-specific and FLAG-specific ZNF143 peaks were lifted over from the hg38 to the hg19 assembly using liftOver. Enrichment was then calculated using the “permTest” function from regioneR^135^ by evaluating the number of overlaps between loop anchors and peaks and using 100 circular permutations to estimate the background. To get the enrichment scores relative to other factors, the enrichments were normalised by dividing all enrichments by the average enrichment, as described in the original study. Proteintech-specific, FLAG-specific, and common peaks were excluded from the mean calculation and the calculated value was retrospectively applied to these datasets to obtain a relative enrichment score. The fraction of loop anchors containing the peaks was calculated using the “intersect” function from BEDTools v2.27.1^127^.

##### CTCF binding to chromatin is independent of ZNF143

A study by Zhou *et al*. previously suggested that a conditional knockout of ZFP143 in haematopoietic stem and progenitor cells affects CTCF binding^38^. The authors reported that in the ZFP143 knockout (KO) condition, CTCF binding was reduced at the majority of CTCF peaks (referred to as ZFP143-related peaks, N=27,801), although some retained the same signal as in the wild-type (WT) condition (referred to as ZFP143-unrelated peaks, N=4196). To investigate the cause of this observation, CTCF ChIP-seq data in the WT and KO cells was re-analysed.

In this study, two independent biological replicates of CTCF ChIP-seq were performed and deposited. Therefore, ChIP-seq CTCF signals from both replicates were used to generate heatmaps over ZFP143-related and ZFP143-unrelated regions. Surprisingly, the loss of signal in the CTCF ChIP-seq dataset over the ZFP143-related peaks could only be detected in the first of the two replicates (Figure S3A). In the second biological replicate, the signal in WT was very similar to the signal in both KO replicates, suggesting potential differences between two CTCF ChIP-seq datasets in the WT, but not KO condition. To understand the nature of this replicate-specific behaviour, orthogonal CTCF ChIP-seq datasets from the same cell type^117,118^ were added. Heatmaps over the same regions revealed that both datasets have very weak CTCF signals over ZFP143-related regions, similar to the CTCF signal from the second WT replicate (Figure S3B). In addition, CTCF motif analysis revealed that CTCF motifs are weak or absent in the ZFP143-related peaks, in contrast to the ZFP143-unrelated peaks, where CTCF motifs have about average strength (Figure S3C). Furthermore, overlapping these two sets of peaks with the compendium of CTCF peaks from CISTROME revealed that only 30% of the ZFP143-related peaks show overlap, whereas this fraction is almost 100% for the ZFP143-unrelated peaks (Figure S3D).

It has been previously reported that ChIP-seq may be subject to GC bias, in some cases leading to a higher proportion of false positive peaks^136^. To investigate whether this might be the cause of the differences between CTCF replicates in WT, the observed versus expected read counts for different GC content bins were calculated. This analysis revealed a severe skew towards a higher than expected number of GC-rich reads in the first replicate of CTCF ChIP-seq in the WT condition, but not in the second replicate and neither of the KO samples, suggesting that an extreme GC bias is present (Figure S3E, S3F).

Taken together, this analysis suggests that the differences in signal in the CTCF ChIP-seq upon ZFP143 knockout are likely a result of GC bias in one of the CTCF ChIP-seq replicates, rather than a true biological change induced by the loss of ZFP143. As only this replicate was used to draw conclusions in the original study, the identified set of ZFP143-related peaks is likely an artefact of the peak calling procedure. The low overlap with a compendium of CTCF peaks and the absence of strong CTCF motifs further suggest that these regions are unlikely to represent actual CTCF binding sites.

##### ZNF143 binding to chromatin is independent of CTCF

Another study by Zhang *et al*. has recently suggested that acute depletion of CTCF in the endometrial carcinoma cell line HEC1B leads to changes in ZNF143 occupancy^39^. The signal in the ZNF143 ChIP-seq dataset was found to be reduced at CTCF binding sites, whereas it was maintained at loci where only ZNF143 signal was present, but not CTCF signal. To investigate this further, we re-analysed the ChIP-nexus signals for ZNF143 and CTCF using the peak coordinates deposited by the authors.

Peaks in the ZNF143 ChIP-nexus dataset from the control HEC1B cells were classified as ZNF143-only (N=15377) or shared (N=19745) based on overlap with CTCF peaks from the same experimental condition using the intersect function of BEDTools v2.27.1^127^. ZNF143 ChIP-nexus signals in control and CTCF-depleted conditions were then used to generate heat maps over the ZNF143-only and shared peaks (Figure S3G, S3H). As reported in the original study, a decrease in ZNF143 signal was observed only for the shared sites. However, the antibody used for ChIP-nexus is the polyclonal anti-ZNF143 Proteintech 16618-1-AP antibody, which we believe to be cross-reacting with CTCF. Therefore, the most parsimonious explanation why ZNF143 signal is lost at CTCF sites is the recognition of CTCF by the Proteintech antibody in the untreated condition and its absence upon CTCF depletion.

This finding suggests that it is unlikely that CTCF positions ZNF143 at CTCF binding sites. The data from Zhou et al suggest that ZNF143 positions CTCF on chromatin, whereas the data from Zhang et al. suggest that CTCF positions ZNF143 on chromatin, albeit at different positions in the genome. Our interpretation of these data is that neither protein affects the positioning of the other, which is in line with our own observation in the acute ZFP143 depletion setting. We would like to emphasise that the independence of CTCF and ZFP143 in chromatin binding and gene regulation is also found in the degron lines from our study and accompanying study (<u>Narducci and Hansen</u>).

##### ZNF143-CTCF motif pairs positioning is an artefact of repetitive elements

Another observation that could indicate a cooperative role for ZFP143 and CTCF chromatin binding is that approximately 60% of ZFP143 motifs were found to be located exactly 37 bp away from the nearest CTCF motif in the mouse genome^38^. ZFP143-CTCF motif pairs provided by the authors were utilised to plot the CTCF and ZFP143 occupancy from the study that reported the motif pairs at these positions. To our surprise, no signal could be detected for either CTCF or ZFP143 in any of the replicates (Figure S3I). To understand whether a specific feature is associated with these motif pairs, a number of genomic characteristics were examined. Strikingly, the reported CTCF-ZFP143 motif pairs completely overlapped with SINE/B2 repetitive elements (Figure S3J). This suggests that the proximity of CTCF and ZFP143 motifs is a chance co-occurrence found in this specific class of repeat elements. The fact that these motif pairs are not bound by either protein makes it unlikely that CTCF or ZFP143 exert any regulatory function at these repeat elements, either in gene regulation or 3D genome organisation.

#### ZFP143 targets

The following procedure was developed to annotate ZFP143-bound genes. All transcripts corresponding to the expressed protein-coding and long non-coding RNA ("transcript_type" field is protein-coding, lincRNA, processed_transcript, bidirectional_promoter_lncRNA or antisense) with high confidence levels ("transcript_support_level" field is 1, 2, 3, 4 or 5) from the GENCODE vM25 gene annotation^137^ were extracted. For each transcript, the coordinates of the TSS window were calculated by taking the coordinates 2 kb upstream and 1 kb downstream of the annotated TSS. The transcripts for which the TSS window overlapped the ZFP143 peaks and the distance between the TSS and the peak centre was less than 2 kb were retained. Next, for each gene, the transcript that met most of the following ranked criteria was retained: highest transcript support level, shortest distance to peak, highest expression and maximum length. Lastly, when the same ZFP143 peak was associated with multiple genes on the same strand, the more appropriate gene was assigned based on (i) transcript type, (ii) transcript support level, and (iii) presence of genes with lower annotation quality at the same location. In particular, protein-coding genes were preferred over lincRNAs, and lincRNAs were preferred over all other types of non-coding RNAs. Genes with transcript support levels of 1 or 2 were prioritised over the others. Finally, genes without canonical names were given a lower priority than any other. If the application of these three rules did not resolve the ambiguous assignment of the ZFP143-bound gene, both genes were discarded.

The described procedure resulted in N=2226 genes being annotated as ZFP143-bound, of which N=1975 were protein-coding genes and N=251 were non-coding RNA. On the other hand, N=1846 ZFP143 peaks were assigned to genes, of which N=375 were located at bidirectionally transcribed promoters. Gene ontology analysis of ZFP143-bound genes was performed using the GO term over-representation test from g:Profiler^138^. The annotation of genes with GO terms from Ensembl (release 110) was utilised, and electronic GO annotations were removed prior to the analysis. An FDR threshold of 0.05 was applied to filter significantly enriched GO terms.

##### Conserved human and mouse ZNF143/ZFP143-bound genes

To analyse the conservation of ZNF143/ZFP143 binding, re-analysed ChIP-seq peaks from human and mouse cell lines were used. All identified peaks for the human and mouse datasets were pooled together and for each peak the number of samples in which the peak was detected was calculated using the “count_overlaps” function from bioframe v0.3.3^139^. The datasets were then classified according to the cell type (32 datasets mapped to 16 cell types in human, 7 datasets mapped to 5 cell types in mouse, Table S2). ZNF143 binding sites were considered conserved across human datasets if a peak was identified in every cell type. The same procedure was followed for mouse ZFP143 datasets. This resulted in N=1456 conserved peaks in human and N=1240 conserved peaks in mouse. 99.5% and 96.2% of the conserved human and mouse peaks, respectively, contained SBS1 and SBS2 motifs, indicating that these are strong ZNF143/ZFP143 binding sites. To assign conserved ZNF143/ZFP143-bound genes, the procedure described above for ZFP143-HA peaks was used, and N=1696 and N=1653 conserved ZNF143/ZFP143-bound genes were annotated in human and mouse, respectively. To map the mouse orthologs of conserved ZNF143-bound human genes, biomaRt^140^ with Ensembl genes (release 110) was used. Gene ontology analysis of the conserved ZNF143/ZFP143-bound genes was performed using the GO term over-representation test from g:Profiler^138^. The annotation of genes with GO terms from Ensembl (release 110) was utilised, and electronic GO annotations were removed prior to the analysis. An FDR threshold of 0.05 was applied to filter significantly enriched GO terms.

#### Hi-C

Hi-C reads were mapped to the mm10 mouse reference genome assembly using bwa mem v0.7.17-r1188^123^ with the Open2C distiller-nf pipeline (https://github.com/open2c/distiller-nf). Mapped reads were parsed using pairtools v0.3.0^141^ with the “walks-policy” parameter set to “mask” and the “max_mismatch_bp” parameter set to 1. Read pairs were then binned into Hi-C contact matrices and written as .cool files. Iterative correction of the matrices and removal of low coverage bins was performed using the "balance" function from cooler v0.8.11^142^.

Genome-wide relative contact probability curves and their derivatives were calculated for the 5 kb resolution Hi-C contact matrices using cooltools v0.5.1^143^. Expected contact matrices were obtained using the "compute-expected" function from cooltools with the "ignore-diags" parameter set to 0. Average loops were calculated for 5 kb resolution observed-over-expected Hi-C contact matrices using coolpup.py v0.9.5^144^ with "pad" set to 200 and "min-dist" set to 0. Loops previously annotated in high-resolution Hi-C and Micro-C datasets^53,54^ were used for the average loop calculations.

#### 4C-seq

4C-seq reads were processed as previously described^56,81^. First, the restriction enzyme (RE) fragment map of the mm10 mouse reference genome was generated for DpnII and Csp6I REs. The fragments derived from mitochondrial DNA and the so-called blind fragments, whose ends were cut by the same restriction enzyme, were removed. Secondly, the sequencing data were demultiplexed per viewpoint by scanning the forward and reverse reads for viewpoint-specific primers, allowing for one mismatch. The forward reads were then trimmed to the start of the DpnII cut site. The resulting reads per viewpoint were mapped to the mm10 mouse reference genome assembly using the bwa aln and bwa samse v0.7.17-r1188^123^. To discard multi-mapping reads, only those with MAPQ > 3 were retained using SAMtools v1.12^124^. Finally, to generate the read counts per RE fragment, for each fragment the number of mapped reads that have the same start or end coordinates as the fragment was counted. To obtain 4C contact profiles, the counts were normalised to the total number of overlaps detected per million reads.

#### TT-seq

TT-seq reads were mapped to the mm10 mouse reference genome assembly using STAR v2.7.9a^145^ with GENCODE vM25 gene annotation^137^. Uniquely mapped reads with MAPQ > 10 were selected using SAMtools v1.12^124^. Duplicate reads were filtered out using the Picard v2.25.6 “MarkDuplicates” function. The reads were split by strand using SAMtools as described previously^52^. Forward strand reads were extracted by including (i) reads that are second in read pair (-f 128), excluding reads that are mapped to the reverse strand (-F 16) and (ii) reads that are first in a pair and are mapped to the reverse strand (-f 80). Reverse strand reads were extracted by including (i) reads that are first in read pair (-f 64), excluding reads that are mapped to the reverse strand (-F 16) and (ii) reads that are second in a pair and are mapped to the reverse strand (-f 144). Gene counts were obtained using the ”htseq-count” function from HTSeq v0.13.5^146^. Counts were calculated separately for genes from forward and reverse strands with the parameters “--stranded no”, “--nonunique all”, “--order pos”, and “--type gene”. Bigwig coverage tracks were generated separately for forward and reverse strands using the deepTools v3.4.2^128^ with the “bamCoverage” function. Scale factors for bigwig generation were calculated as the number of reads per sample adjusted to match the coverage of the sample with the highest coverage. Genomic tracks plots were produced using pyGenomeTracks v3.8^131^.

Differentially expressed genes were identified using the DESeq2 v1.30.1^147^. Low expressed genes were filtered by requiring half of the samples to have gene counts greater than 10. The “nbinomWaldTest” function with default parameters was used to test contrasts of dTAG-V1 treatment time against DMSO. MitoCarta v3.0^62^ was used to annotate the differentially expressed genes with the mitochondrial localisation, pathways, and processes. Gene set enrichment analysis was performed using GSEA v4.3.2^148^ in pre-ranked mode with 10,000 permutations and gene sets from MSigDB v2023.2^149^. Gene sets with FDR < 0.05 were considered significant.

#### Mass-spectrometry

Raw data were analysed by DIA-NN v1.8^150^ without a spectral library and with the “Deep learning” option enabled. The Swissprot Mus Musculus database (17.125 entries, release 2022_08) was added for the library-free search. The quantification strategy was set to Robust LC (high accuracy) and the MBR option was enabled. The other settings were kept at the default values. The protein groups report from DIA-NN was used for downstream analysis in Perseus v2.0.10.0^151^. Values were log2-transformed, after which proteins were filtered for at least 100% valid values in at least one sample group. Missing values were replaced by imputation based on a normal distribution with a width of 0.3 and a minimal downshift of 2.4. Differentially expressed proteins were determined using a Student’s t-test (minimal threshold: -log(p-value) ≥ 1.3 and [x-y] ≥ 0.5 | [x-y] ≤ -0.5).

Gene set enrichment analysis was performed using GSEA v4.3.2^148^ in pre-ranked mode with 10,000 permutations and gene sets from MSigDB v2023.2^149^. Genes corresponding to the identified proteins were pre-ranked by the protein abundance difference derived from differential expression analysis. Gene sets with FDR < 0.05 were considered significant.

#### RNA-seq

RNA-seq reads were mapped to the mm10 mouse reference genome assembly using STAR v2.7.9a^145^ with GENCODE vM25 gene annotation^137^. Read counts per gene were obtained using the "quantMode" parameter in STAR. Differentially expressed genes were identified using the DESeq2 v1.30.1^147^. Low expressed genes were filtered by requiring half of the samples to have gene counts greater than 10. The “nbinomWaldTest” function with default parameters was used to test contrasts. Test results at the lfcThreshold of log2(1.5) were extracted and used for downstream analyses. Distances between the samples were calculated based on the VST-transformed gene count matrix using the “plotPCA” function. Gene set enrichment analysis was performed using GSEA v4.3.2^148^ in pre-ranked mode with 10,000 permutations and gene sets from MSigDB v2023.2^149^. Gene sets with FDR < 0.1 were considered significant.

#### ATAC-seq

ATAC-seq reads were mapped to the mm10 mouse reference genome assembly bwa mem v0.7.17-r1188^123^. Uniquely mapped reads that were not mapped to mitochondrial DNA and had MAPQ > 10 were selected using SAMtools v1.12^124^. Duplicate reads were filtered out using the Picard v2.25.6 “MarkDuplicates” function.

To deconvolve cell type compositions in bulk ATAC-seq samples from control and ZFP143-depleted gastruloid, previously published single-cell ATAC-seq atlas of gastruloid development was used^74^. Firstly, pseudobulk ATAC-seq profiles for each cell type (cluster) were generated from single-cell data by summing counts in each peak across all cells belonging to the same clusters. Then, the read counts of DMSO and dTAG-V1 treated bulk samples in the same peak set used in the single-cell data were generated. Bulk and pseudobulk counts were concatenated, normalised using the median of ratios normalisation implemented in DESeq2 v1.30.1^147^, and transformed using the variance stabilizing transform (VST).

Since bulk samples can be thought of as linear combinations of different cell types, independent component analysis (ICA) can be used to unmix the contributions of constituent cell types in bulk samples. ICA is a method for separating a multivariate signal into a predefined number of additive, mutually independent components. Briefly, ICA was performed on VST-transformed and zero-centred pseudobulk counts, using the FastICA algorithm, implemented in the “icafast” function from ica R package, with number of components set to 7 and a maximum of 100 algorithm iterations. The source signal estimates and the unmixing matrix obtained from ICA were subsequently used on the VST-transformed bulk count matrix to unmix the cell type contributions. Finally, the cell type proportion estimates were obtained by setting the negative contributions to zero and normalising contributions to the sum of the contributions for each sample.

#### Image analysis

Gastruloid brightfield images were analysed using ImageJ v2.9.0^152^ to quantify morphological features. Files were opened using Bio-formats^153^. Images were pre-processed by applying a Gaussian filter (sigma=3) to smooth the images and then the local contrast was normalised (x=200, y=200, STD=3, centered). The gastruloid outline was selected using a supervised thresholding, converted into a binary mask, and consequently analysed with the “Analyse particles” plugin. Briefly, the gastruloid outline is selected as a region of interest (ROI) and shape descriptors, such as “Roundness”, “Aspect ratio”, and “Area” were calculated.

### Key resources table

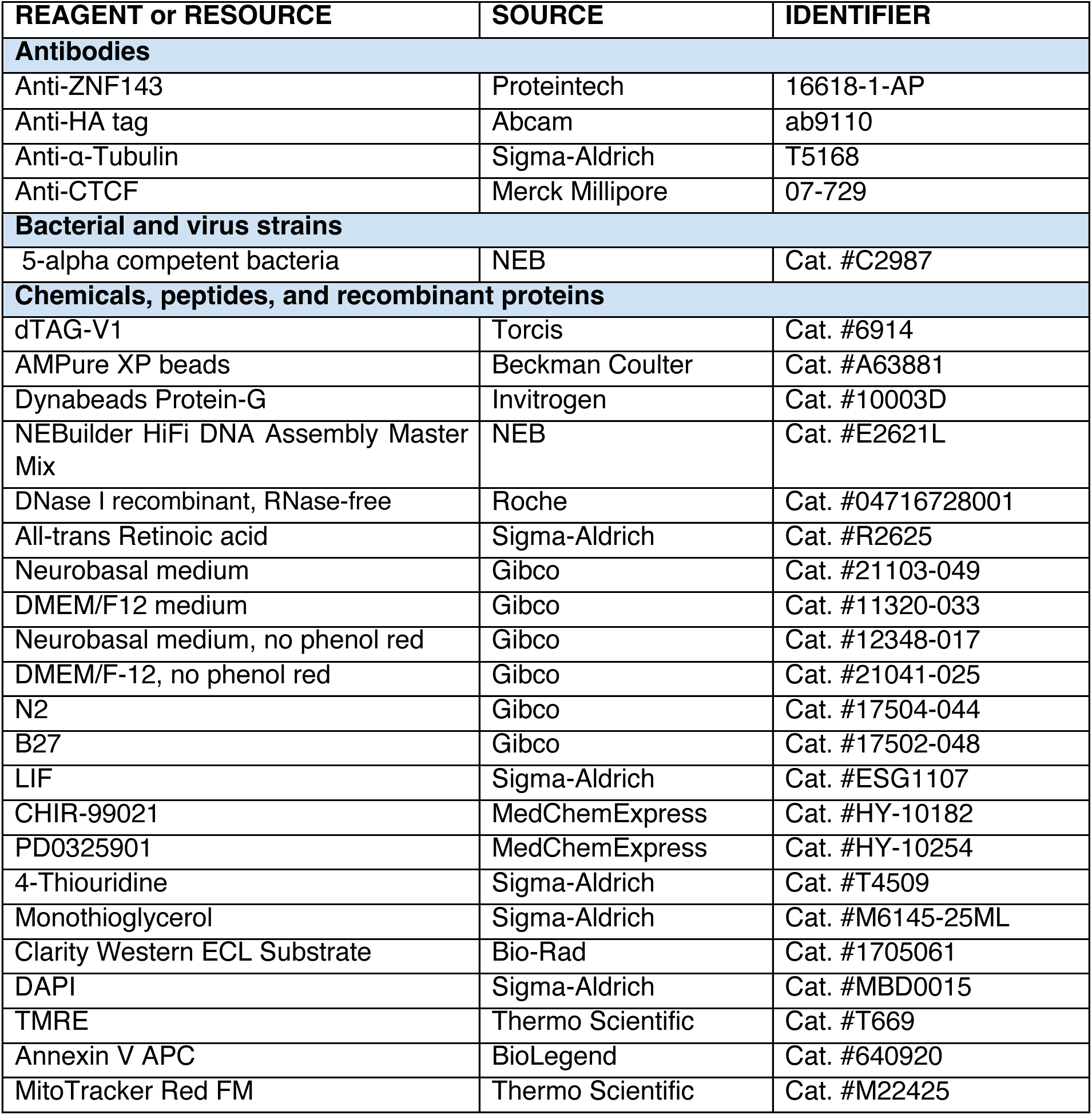
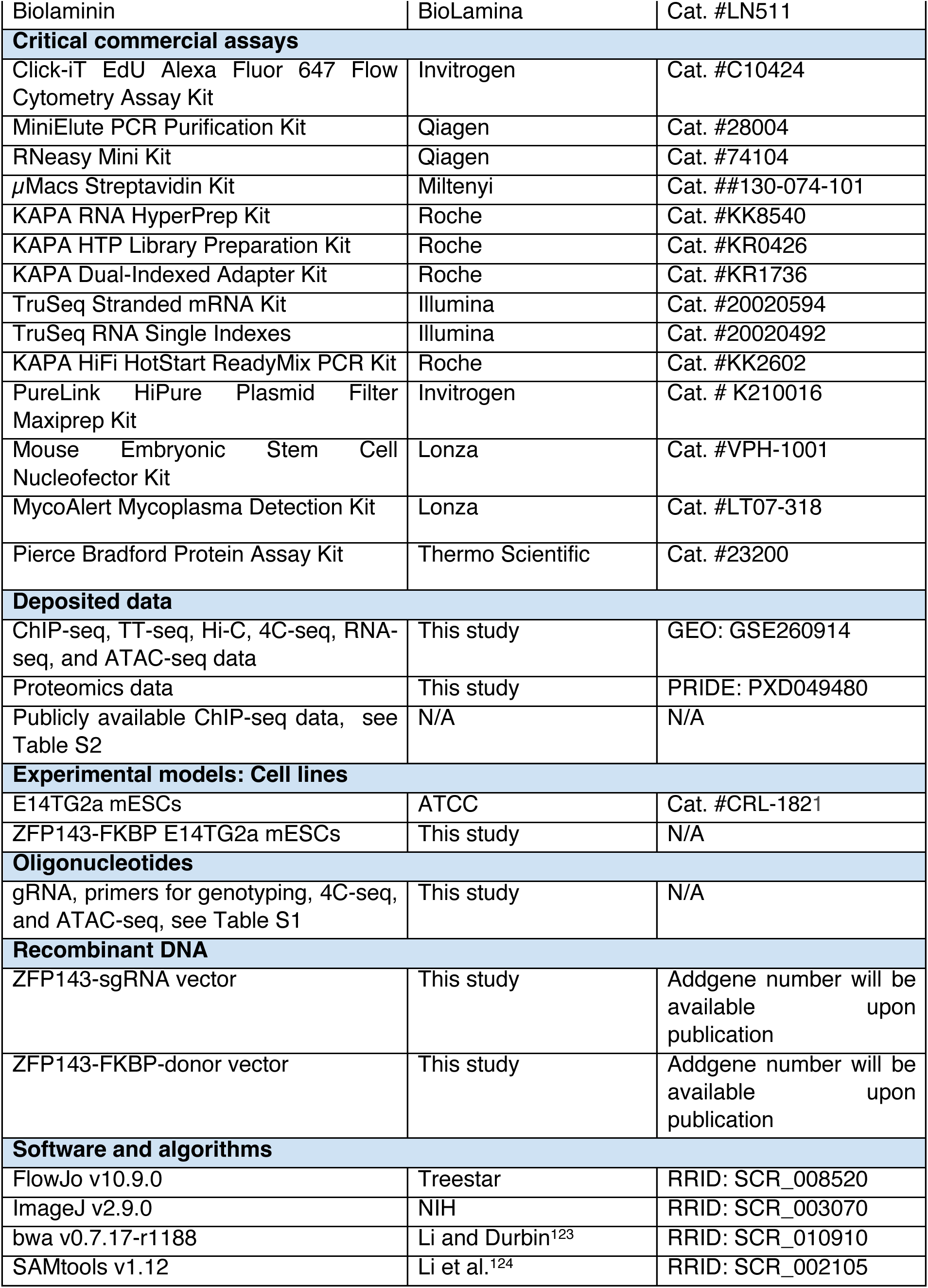
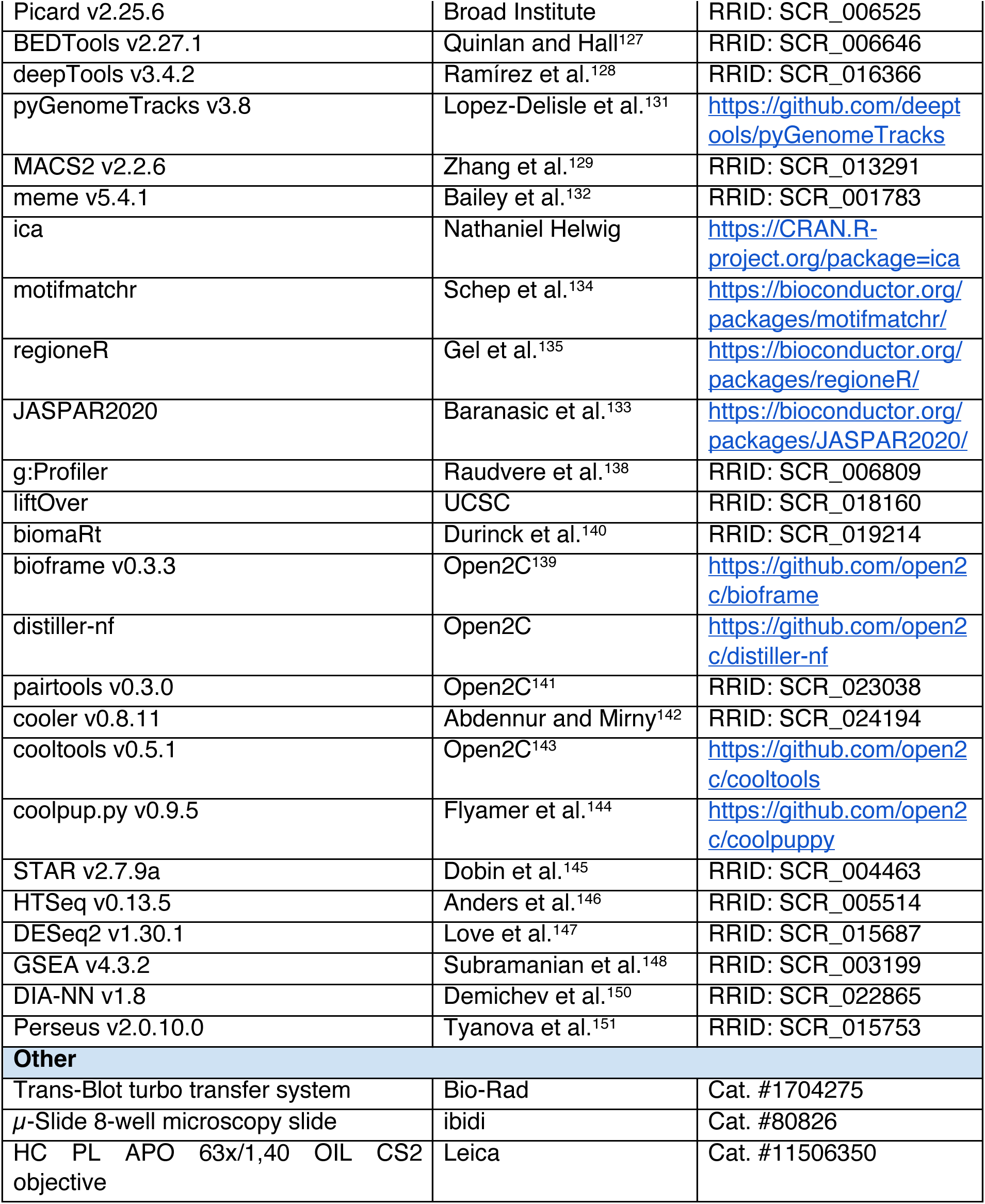

